# Matrix biophysical cues drive transient or prolonged breast cancer dormancy programmes

**DOI:** 10.64898/2026.10.28.762410

**Authors:** Unai Heras, José Miguel Pardo-Sánchez, Jone Berasain, Iune Toledo, Leire Iturriaga, Itxaso Calafel, Robert Aguirresarobe, Guillermo de Cárcer, Julio A. Aguirre-Ghiso, Elena Arizaga-Batiz, Marta Fernandez-Calleja, Rosa M. Zubizarreta-Susperregui, Ana Legorburu, Ana Riestra, Oihane Mitxelena-Iribarren, Amaia Cipitria

## Abstract

Breast cancer metastasis can emerge after disease-free periods through the reawakening of dormant disseminated cancer cells. Despite evidence that the extracellular matrix (ECM) modulates this phenomenon, biophysical cues regulating dormancy fate and therapeutic responses remain poorly understood. Here, we engineered stiffness-matched gelatin methacryloyl (GelMA) and alginate-based 3D matrices to disentangle matrix stiffness from adhesion and degradability and uncover their contributions to dormancy. Deep learning-based 3D segmentation revealed that adhesive and degradable matrices supported rounded and proliferative tumour masses, whereas inert matrices favoured single cell dormancy. Within inert matrices, stiffness further determined dormancy fate, with soft matrices activating SOX9-associated transient stem-like dormancy, while stiff matrices reduced cell volume, increased intracellular macromolecular crowding of proteins and induced NR2F1-associated prolonged dormancy marked by autophagy and cellular stress. Transiently dormant cells retrieved from soft matrices retained dormancy memory and failed to form tumours in chicken chorioallantoic membranes, demonstrating persistence of matrix-induced states after removal of biophysical cues. Moreover, patient-derived cells in soft inert matrices recapitulated transient dormant states and exhibited therapy resistance. We demonstrate that tuneable biophysical cues shape distinct dormancy states, linking matrix mechanics to dormancy regulation and therapeutic response.

## INTRODUCTION

Metastasis is the leading cause of breast cancer (BC) mortality, accounting for 80 % of cancer-related deaths ^1,2^. Disseminated cancer cells (DCCs) are detected in the bone marrow of approximately 27 % of early-stage BC patients, associated with late recurrence and poor clinical outcome ^3,4^. Luminal BC shows a strong tropism for bone, where DCCs persist in a clinically undetectable dormant state for years before reawakening ^5–9^. However, the inability to detect single dormant DCCs by imaging technologies or to routinely collect bone marrow aspirations from patients ^10^ limits mechanistic studies and therapeutic strategies.

Dormancy has been extensively investigated using genetically engineered mouse models ^11,12^, establishing that both extrinsic (i.e. extracellular matrix (ECM)) and intrinsic (transcriptional and epigenetic) mechanisms regulate dormancy ^7,13^. ECM stiffness ^14^, composition ^15^ and architecture ^16^ are sensed by DCCs and integrated into dormancy-associated transcriptional programmes that determine dormancy or reactivation. For example, SOX-family transcription factors influence stemness, epithelial-to-mesenchymal (EMT) programmes, and endocrine therapy resistance, while NR2F1 acts as a barrier for early dissemination, and in DCCs that eventually lodge in target organs, it promotes prolonged dormancy ^17–21^. Yet, how specific ECM biophysical cues in target organs are translated into dormancy-associated transcription programmes remains poorly understood ^7^, as they are difficult to decouple *in vivo*.

3D bioengineered matrices provide controlled systems to investigate how cell-matrix interactions regulate dormancy ^22–27^ by independently tuning matrix biophysical ^26–32^ and biochemical ^24,33–35^ cues. Nevertheless, most have primarily focused on tumour mass dormancy and the contribution of specific biophysical cues on growth-arrested single cells remains poorly resolved ^5,36,37^. Moreover, they rarely incorporate patient-derived cells ^29,31,38–41^, limiting the understanding whether biophysical cues regulate therapeutic responses beyond established cell lines ^42^.

Here, we engineered stiffness-matched gelatin methacryloyl (GelMA) and alginate-based hydrogels to disentangle the effects of matrix stiffness from adhesion and degradability and determine how these properties regulate BC dormancy fate. Using luminal BC MCF-7 and patient-derived cells, we quantified cell-cycle dynamics, 3D volumetric and morphometric changes of single cells versus tumour masses using deep learning-based 3D segmentation models, identified stiffness-dependent transient or prolonged dormancy states associated with macromolecular crowding of proteins and SOX9 and NR2F1 transcription factors, evaluated the persistence of matrix-induced dormancy states after removal of the biophysical cues through tumour formation capacity in chicken chorioallantoic membrane (CAM) models, and recapitulated dormant states and their response to endocrine therapy in patient-derived cells.

## RESULTS

### Stiffness-matched hydrogels reveal opposing cell-cycle dynamics

Given that bone marrow stiffness ranges from 0.25-25 kPa ^43,44^, following previous work ^29^, we tuned 3D hydrogel stiffness to 1 and 10 kPa to model soft and stiff bone marrow compartments, respectively (Fig. 1a-c).

**Fig. 1.**
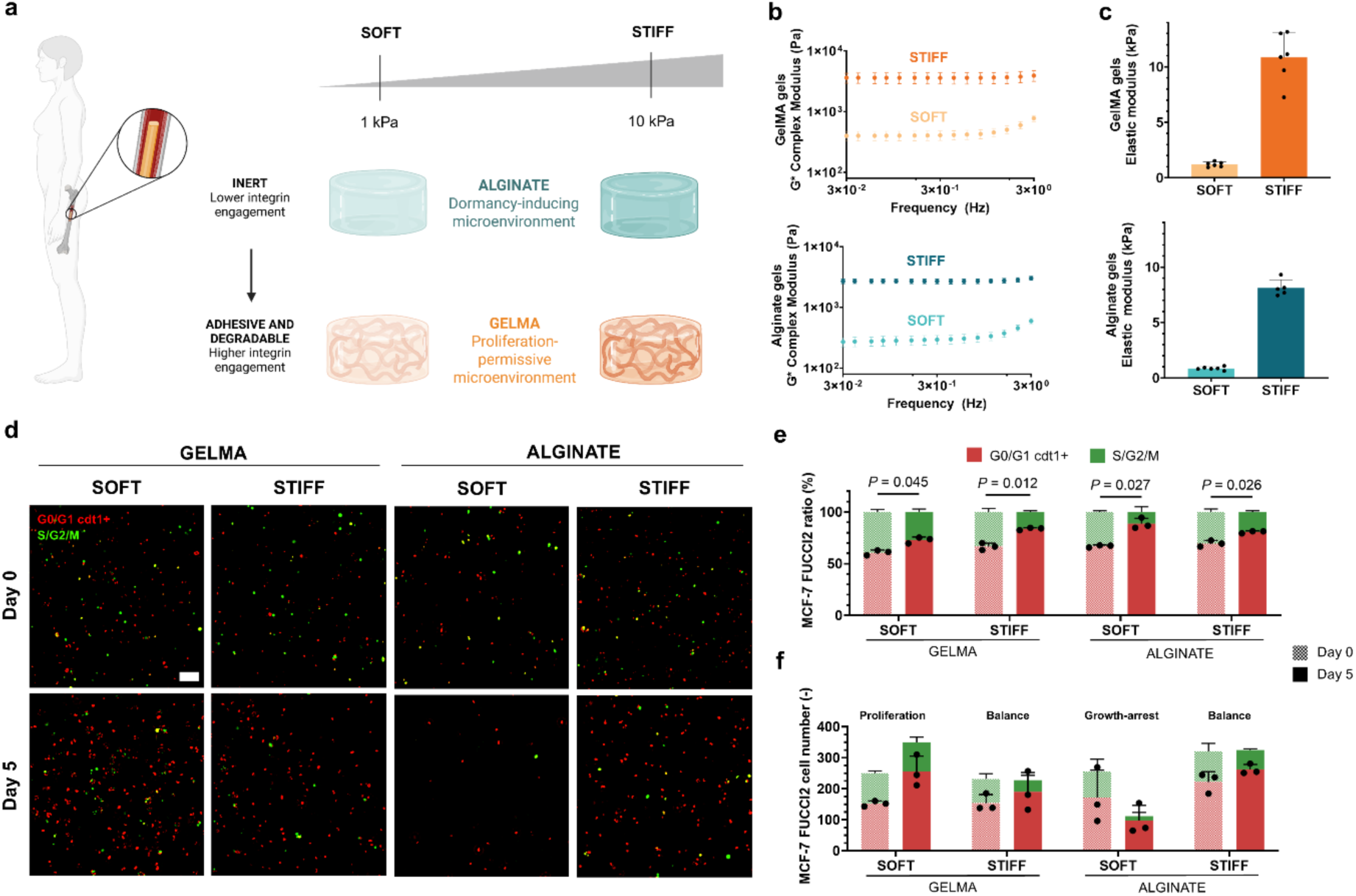
Stiffness-matched 3D hydrogels with distinct adhesion and degradability cues reveal opposing cell-cycle states. **a**, Graphical illustration of 3D hydrogels mimicking bone marrow stiffness. Adhesive and degradable GelMA represents a proliferation-permissive microenvironment, whereas inert, non-adhesive and non-degradable alginate represents a dormancy-inducing microenvironment. Schematics created with BioRender. **b**, Rheological measurements of GelMA and alginate hydrogels showing complex modulus (G*) values for soft and stiff hydrogels at day 5 with MCF-7 cells (n = 6). **c**, Corresponding elastic moduli (n = 6). **d**, Representative maximum projection of MCF-7 FUCCI2 images showing the cell cycle reporter (G_0_/G_1_ cdt1^+^ = mCherry-hCdt1 and S/G_2_/M = mVenus-hGeminin) in soft and stiff GelMA and alginate at day 0 and 5 (200 μm z-stacks, 10x objective, n = 3). Scale bar, 100 μm. **e**, Cell cycle ratio distribution and **f**, absolute numbers of G_0_/G_1_ cdt1^+^ and S/G_2_/M at day 0 and 5 (n = 3). Bars and error bars show mean ± s.d. Statistical significance (two-tailed Student’s t-test (parametric)).

Stiffness was matched and decoupled by comparing adhesive and degradable GelMA with adjustable polymer concentration to inert norbornene-modified alginate hydrogels with varying dithiothreitol (DTT) crosslinker concentration. Formulations were adjusted according to the actual degree of substitution (DS) of each biomaterial (83% for GelMA and 21.58 % for alginate) and validated by stiffness and mesh size values without cells (Supplementary Fig. 1).

MCF-7 viability was evaluated by the number of live cell units, live/dead ratio and live unit area (Supplementary Fig. 2a-d). At day 5, cell viability was preserved at above 150 live cell units in both GelMA and alginate hydrogels, across soft and stiff conditions. By day 14, high viability ratio was preserved above 80% in GelMA hydrogels, with fewer live cell units but an increase in live cell area. In contrast, cell viability decreased to 25-40% in alginate, with a decrease in both live units and live unit area. Adenosine triphosphate (ATP) release confirmed these findings, with higher metabolic activity in soft and stiff GelMA than in soft and stiff alginate (Supplementary Fig. 2e,f). Since viability (ratio above 50%) was highest at day 5, subsequent analyses focused on this time point.

To assess proliferative versus growth-arrested behaviours, we encapsulated MCF-7 cells with fluorescence-ubiquitination-based cell cycle indicator 2 (FUCCI2) and analysed the cell cycle distribution ^45,46^. FUCCI2 analysis revealed a significantly higher proportion of G0/G1 cdt1^+^ cells across all conditions (Fig. 1d,e). Absolute numbers indicated that cells in stiff GelMA and alginate displayed a balanced survivor behaviour (Fig. 1f).

In contrast, stiffness-matched soft hydrogels revealed that G0/G1 cdt1^+^ cells increased in GelMA (indicative of proliferation), whereas both G0/G1 cdt1^+^ and S/G2/M cell populations decreased in soft alginate, with loss of FUCCI2 fluorescence, in line with low cdt1 protein expression (G0/G1 cdt1^-^) indicating growth-arrest ^47^, as previously reported in 3D alginate studies ^29^.

### Biophysical cues shape 3D single cell and tumour mass volume and morphology

We quantified collective populations of single cells and tumour masses (clusters with ≥ 2 nuclei) across hydrogels over time using deep learning models for 3D segmentation (Supplementary Fig. 3). Proliferation-permissive GelMA generated more tumour masses at day 5, while inert alginate retained predominantly single cells, with significant differences only in soft matrices (Fig. 2a,b). By day 14, population distributions remained stable (Supplementary Fig. 4a-c).

**Fig. 2.**
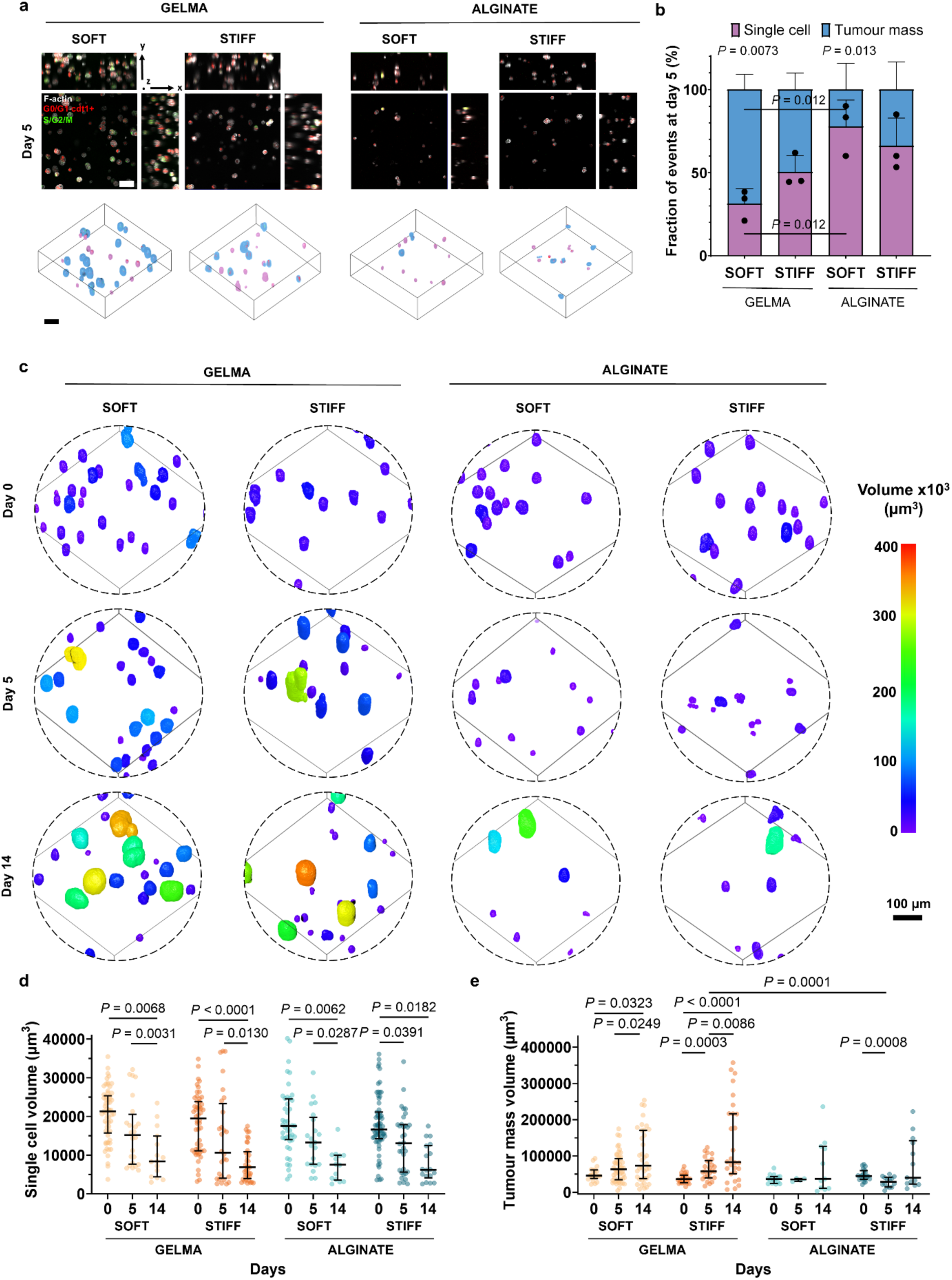
3D volumetric analysis in collective populations of single cells and tumour masses. **a**, Representative confocal MCF-7 FUCCI2 and F-actin staining in GelMA and alginate hydrogels at day 5 (200 μm z-stacks, 20x objective, n = 3). 3D reconstruction in ORS Dragonfly where blue represents tumour masses and purple refers to single cells. Scale bar, 100 μm. **b**, Fraction of single cells versus tumour masses at day 5 (n = 3, mean ± s.d.). **c**, 3D reconstruction and segmentation in ORS Dragonfly of confocal MCF-7 FUCCI2 and F-actin staining at days 0, 5 and 14 in soft and stiff GelMA and alginate hydrogels (200 μm z-stacks, 20x objective, n = 16 - 67 units from n = 3 hydrogels per group). **d**, Single cell and **e**, tumour mass volume quantification at days 0, 5 and 14. Lines and error bars show median and interquartile range, respectively. Statistical significance (two-tailed Student’s t-test (parametric) and Mann-Whitney U-test (non-parametric)).

Next, we investigated 3D volumetric changes. At day 5, stiff alginate reduced the volume of both single cells and tumour masses (Fig. 2c-e), while stiff GelMA increased tumour mass volume (Fig. 2e). By day 14, surviving single cells significantly reduced in volume across all conditions, while tumour masses in both GelMA hydrogels increased in volume.

We characterised epithelial- and mesenchymal-like states associated to dormancy ^48–50^ by E-cadherin and Vimentin localisation within soft groups, respectively, where significant differences in their distribution were observed. In soft GelMA, single cells and tumour masses maintained an epithelial-like phenotype at day 5, with stable membrane E-cadherin localisation (Fig. 3a,b). In contrast, in soft alginate E-cadherin localisation shifted from the membrane to the perinuclear and nuclear regions by day 5, with no changes in Vimentin expression and localisation (Fig. 3a-c). Stiff alginate counterpart showed no changes in E-cadherin localisation (Supplementary Fig. 4d,e).

**Fig. 3.**
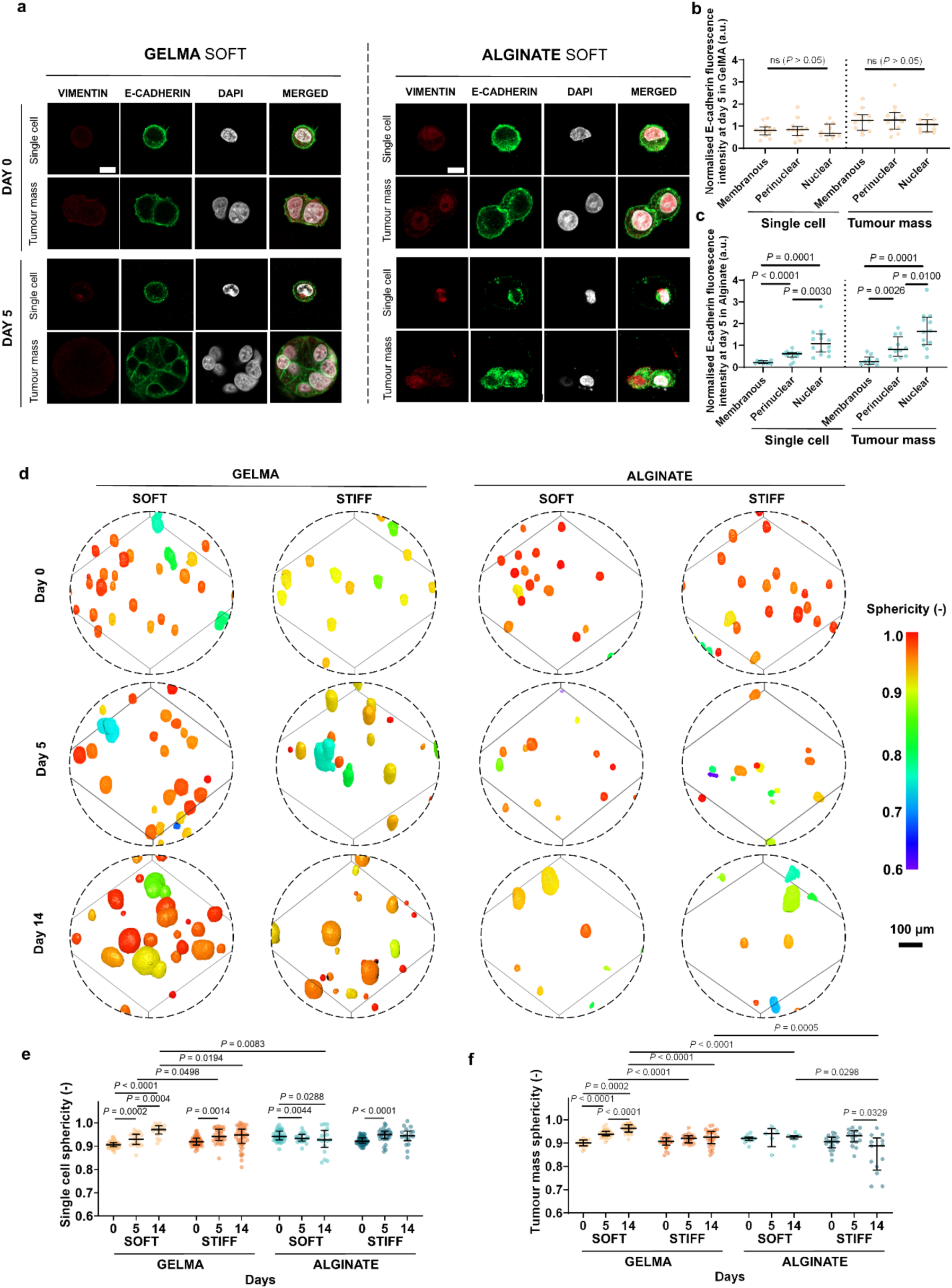
Mesenchymal versus epithelial characterisation in soft hydrogels and 3D morphometric analysis of collective populations of single cells and tumour masses. **a**, Representative confocal 2D images of MCF-7 cells stained for Vimentin (red) and E-cadherin (green) at day 0 and 5 in soft GelMA and alginate hydrogels. Cells are classified as single cells and tumour masses (DAPI = white). Scale bars, 10 μm. **b-c**, Quantification of E-cadherin intensity in membranous, perinuclear and nuclear regions at day 5, normalised to day 0 in (b) soft GelMA and (c) alginate (n = 12 units from n = 3 hydrogels per group). **d**, 3D reconstruction and segmentation in ORS Dragonfly of confocal MCF-7 FUCCI2 and F-actin staining at days 0, 5 and 14 in soft and stiff GelMA and alginate hydrogels (200 μm z-stacks, 20x objective, n = 16-67 unit from n = 3 hydrogels per group). **e**, Single cell and **f**, tumour mass sphericity quantification at days 0, 5 and 14. Lines and error bars show median and interquartile range, respectively. Statistical significance (two-tailed Student’s t-test (parametric) and Mann-Whitney U-test (non-parametric)). a.u. stands for arbitrary units.

We further characterised 3D morphometric changes through sphericity analyses. At day 5, single cells adopted a more rounded morphology in all groups, except in soft alginate, where they became more elongated (Fig. 3d,e). By day 14, these trends were enhanced in soft groups, while cells in the stiff groups maintained a stable morphology. Among tumour masses, only those in soft GelMA exhibited a more rounded morphology at day 5 (Fig. 3f), different from tumour masses in stiff GelMA. Tumour masses in stiff alginate became elongated by day 14.

Finally, we assessed nuclear volume over time (Supplementary Fig. 5a). By day 5, only single cells within stiff groups showed larger nuclei (Supplementary Fig. 5b). Tumour masses in soft alginate and stiff GelMA also showed enlarged nuclear volumes by day 5 (Supplementary Fig. 5c). These differences disappeared by day 14. In contrast, tumour masses in soft GelMA showed a progressive decrease in nuclear volume, consistent with previous reports of G0/G1 cdt1+ cell accumulation in 3D hydrogels ^51^. Stiff GelMA tumour masses contained the highest number of nuclei at day 14 (median of 10 nuclei per cluster) compared to the other groups (3-5 nuclei per cluster) (Supplementary Fig. 5d).

### Biophysical cues regulate proliferative versus dormant phenotypes

We characterised proliferative and dormant phenotypes of collective populations of single cells and tumour masses by Ki67 (proliferation), p27 (dormancy), and the phosphorylated forms of ERK (proliferation) and p38 (stress) through the P-ERK:P-p38 ratio (low for dormancy). Given the early morphological changes and high viability observed on day 5, we focused the subsequent studies on that timepoint.

We observed approximately 50% Ki67-positive (Ki67^+^) cells in soft and stiff GelMA, whereas most cells in soft and stiff alginate remained Ki67-negative (Ki67^-^) (Fig. 4a,b). Single cells lacked Ki67 nuclear expression across all conditions (Fig. 4c). Only tumour masses in soft and stiff GelMA displayed Ki67 nuclear expression. Tumour masses in soft and stiff alginate remained Ki67^-^, indicating a non-proliferative phenotype.

**Fig. 4.**
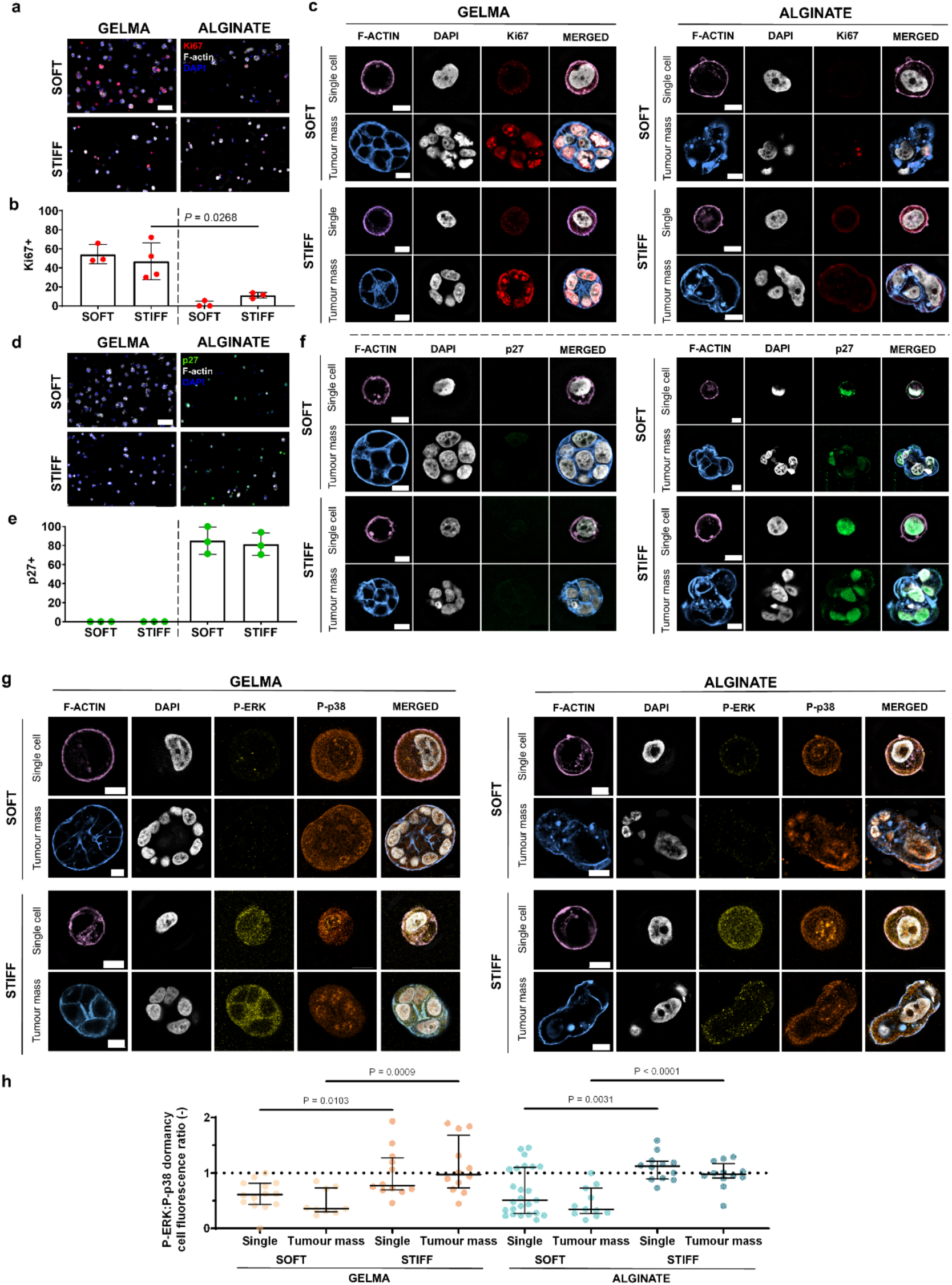
Dormancy characterisation of collective populations of single cells and tumour masses through proliferation and dormancy markers. **a**, Representative maximum projection fluorescence images of MCF-7 cells stained for Ki67 (red) in soft and stiff GelMA and alginate hydrogels at day 5 of experiment (F-actin = white, DAPI = blue) (200 μm z-stacks, 20x objective, n = 3, mean ± s.d.). Scale bar, 100 μm. **b**, Fraction of Ki67^+^ cells across all groups (n = 5 - 43 units from n = 3 hydrogels per group, mean ± s.d.). **c**, Representative confocal 2D images of MCF-7 cells stained for Ki67 (red) in single cells and tumour masses at day 5. Scale bar, 10 μm. **d**, Representative maximum projection fluorescence images of p27 (green) under the same conditions (200 μm z-stacks, 20x objective, n = 3). Scale bar, 100 μm. **e**, Fraction of p27^+^ cells across all groups (n = 10 - 35 units from n = 3 hydrogels per group, mean ± s.d.). **f**, Representative confocal 2D images of p27 (green) in single cells and tumour masses under the same conditions. Scale bar, 10 μm **g**, Representative confocal MCF-7 cell images stained with P-ERK (yellow) and P-p38 (orange) soft and stiff GelMA and alginate hydrogels at day 5. Cells are classified as single cells and tumour masses (DAPI = white). Scale bars, 10 μm. **h**, P-ERK:P-p38 fluorescence intensity ratio (n = 9 - 25 units from n = 3 hydrogels per group). Lines and error bars show median and interquartile range, respectively. Statistical significance (two-tailed Student’s t-test (parametric) and Mann-Whitney U-test (non-parametric)).

We validated these findings by p27 expression. In soft and stiff GelMA no detectable p27-positive (p27^+^) cells were observed, whereas in soft and stiff alginate approximately 80% of the cells were p27^+^ (Fig. 4d,e). Single cells and tumour masses in alginate showed consistent nuclear p27 expression compared to GelMA (Fig. 4f).

Having unravelled the role of adhesion and degradability by comparing GelMA and alginate hydrogels, we next examined whether matrix stiffness modulates P-ERK:P-p38 stress-associated dormancy ^29,52^. P-ERK:P-p38 ratio was significantly lower in both single cells and tumour masses in soft compared to stiff groups (Fig. 4g,h), indicating stress-associated signalling of MCF-7 cells in soft groups. However, this ratio alone did not define dormancy as it was similarly reduced in soft GelMA. Therefore, subsequent analyses focused on p27^+^ dormant cells in alginate hydrogels.

### Matrix stiffness determines distinct transcriptional dormancy states

To investigate how stiffness modulates gene expression in luminal BC dormancy, we compared RNA-sequencing (RNA-seq) datasets of cells retrieved from soft and stiff alginate at day 5 (Supplementary Fig. 6). Gene ontology (GO) enrichment of differentially expressed genes (DEGs) identified protein localisation, cell cycle phase transitions, autophagy, endoplasmic reticulum stress and unfolded protein response (UPR) as the dominant biological processes (Fig. 5a). Distinct gene set signatures further differed between both groups (Fig 5b), indicating enrichment of zinc finger protein genes (*Znf827, Znf804b*) in soft alginate and cyclin-dependent kinase (CDK) inhibitor p57 (*Cdkn1c*) in stiff alginate (Fig. 5c), among others.

**Fig. 5.**
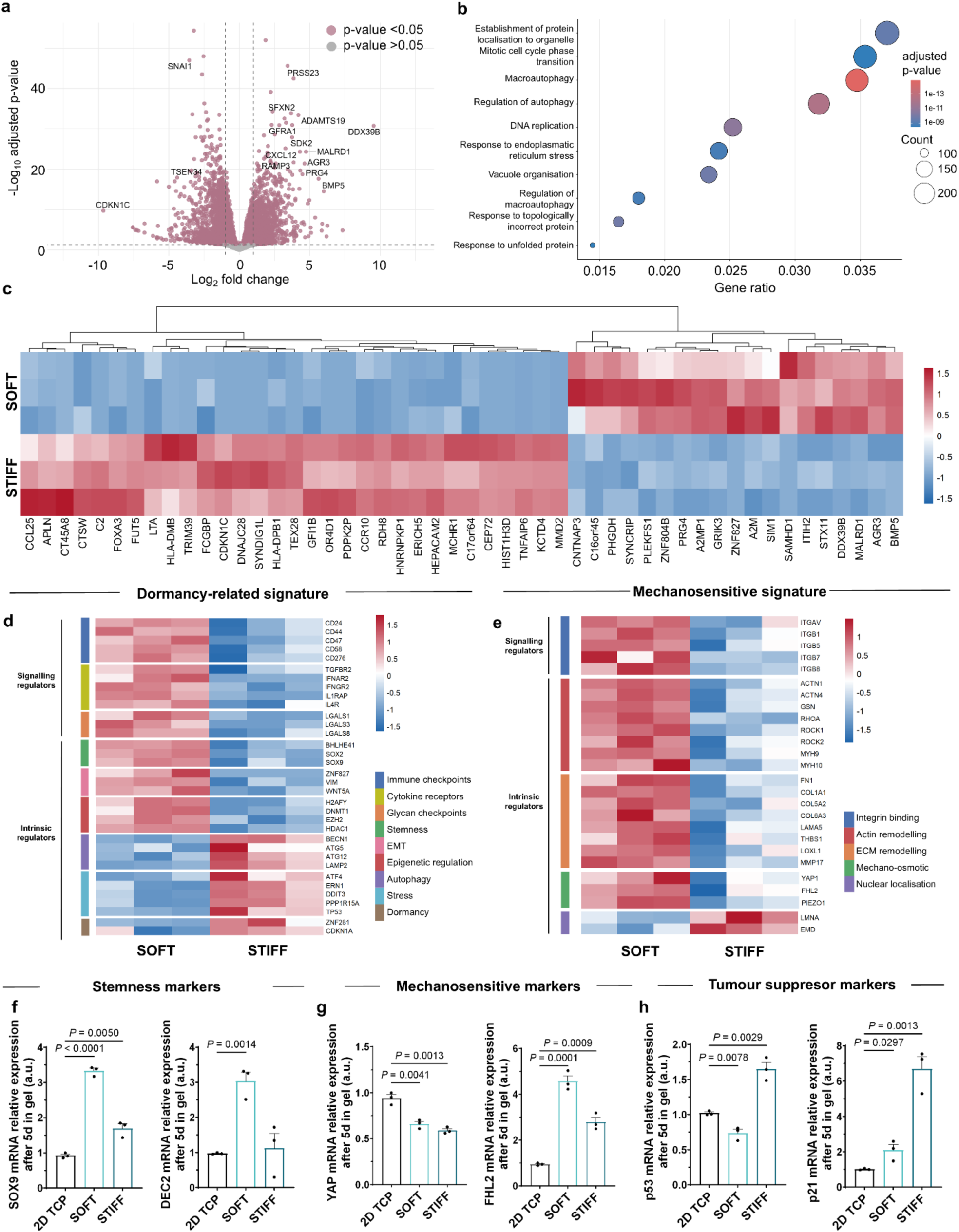
Gene expression analysis in MCF-7 cells in soft versus stiff alginate. **a**, Volcano plot of DEGs between soft and stiff alginate (grey = not significant, salmon = adjusted P-value < 0.05, n = 15172 genes). **b**, GO biological process enrichment of DEGs (adjusted P-value < 0.1) identified by over-representation analysis. **c**, Heatmap of the top 50 DEGs ranked by absolute log2 fold change (variance-stabilised transformed; VST). **d-e**, Expression heatmaps of selected candidate genes associated with (d) potential dormancy-related signature and (e) mechanosensitive signature, classified into signalling and intrinsic regulators. **f-h**, RT-qPCR validation of selected transcripts in MCF-7 cells encapsulated within soft and stiff alginate and retrieved after 5 days (n = 3 biological replicates), compared to 2D controls. Bars and error bars show mean ± s.d., respectively. Statistical significance (two-tailed Student’s t-test (parametric)). a.u. stands for arbitrary units.

We next analysed a gene signature (Fig. 5d). Soft alginate preferentially upregulated signalling regulatory genes, including immune checkpoint genes (*Cd24, Cd44, Cd47, Cd58, Cd276*), cytokine receptors (*Tgfbr2, Ifnar2, Ifngr2, Il1rap, Il4r*), and galectins (*Lgals1, Lgals3, Lgals8*), previously linked to an immunosuppressive phenotype. Soft alginate also enriched for genes associated with dormancy, stemness and plasticity (*Bhlhe41, Sox2, Sox9*), EMT (*Znf827, Vim, Wnt5a*) and epigenetic regulation (*H2afy, Dnmt1, Ezh2, Hdac1*). In contrast, stiff alginate upregulated genes associated with autophagy (*Becn1, Atg5, Atg12, Lamp2*), UPR (*Atf4, Ern1, Ddit3, Ppp1r15a*), tumour suppression (*Tp53*), CDK inhibitor p21 (*Cdkn1a*) and long-term dormancy (*Znf281*). Overall, these data suggest that soft matrices promote stem-like growth arrest phenotypes with some genes expressed in models of dormancy and human samples ^53^, whereas stiff matrices favoured autophagic and stress-adaptative phenotypes associated also with dormancy genes found in prolonged dormancy models.

We also analysed a mechanosensitive signature (Fig. 5e). Soft alginate upregulated integrin genes (*Itgav, Itgb5, Itgb7, Itgb8*), including β1 integrin (*Itgb1*), a key component of fibronectin-binding receptor α5β1. Mechanosensitive and osmotic regulators (*Yap1, Fhl2, Piezo1*), actin cytoskeletal contractility (*Actn1, Gsn, Rhoa, Rock1,* etc.) and ECM remodelling genes (*Fn1, Col1a1, Thbs1, Loxl1,* etc.) were also upregulated in soft alginate, consistent with morphological changes, stemness and EMT traits. Conversely, stiff alginate upregulated nuclear structural genes (*Lmna, Emd*).

Reverse transcription quantitative PCR (RT-qPCR) validated upregulation of *Sox9*, dec2 (*Bhlhe41*) and mechanosensitive *Fhl2* genes in soft alginate (Fig. 5f,g), as well as tumour suppressor *Tp53* and p21 (*Cdkn1a*) genes in stiff alginate (Fig. 5h). Compared with proliferative 2D tissue-culture plates (TCP), most dormancy-associated genes were significantly upregulated in 3D alginate. Notably, *Tp53* was downregulated in soft alginate, suggesting a *Tp53-*independent stem-like dormant phenotype. *Yap1* was also reduced in both soft and stiff alginate compared to 2D TCP, indicating suppression of mechanosensitive signalling in dormant MCF-7 cells.

### Stiffness modulates macromolecular crowding and dormancy-associated transcription factors

We examined intracellular macromolecular crowding of proteins by using the CRONOS FRET sensor ^54,55^. This biosensor consists of an mNeonGreen (mNG) donor and an mScarlet-I (mScal) acceptor fluorophore, connected by a conformationally flexible linker composed of two α-helical peptides joined by a hinge.

Under high intracellular macromolecular crowding conditions, the sensor conformation becomes more compact, resulting in increased FRET efficiency (Fig. 6a). MCF-7 cells expressing CRONOS sensor exhibited significantly higher FRET signal in stiff alginate compared to the soft group (Fig. 6b,c), also normalised to the donor fluorophore expression (Fig. 6d). This indicates elevated intracellular macromolecular crowding in prolonged dormancy programs induced by stiff alginate.

**Fig. 6.**
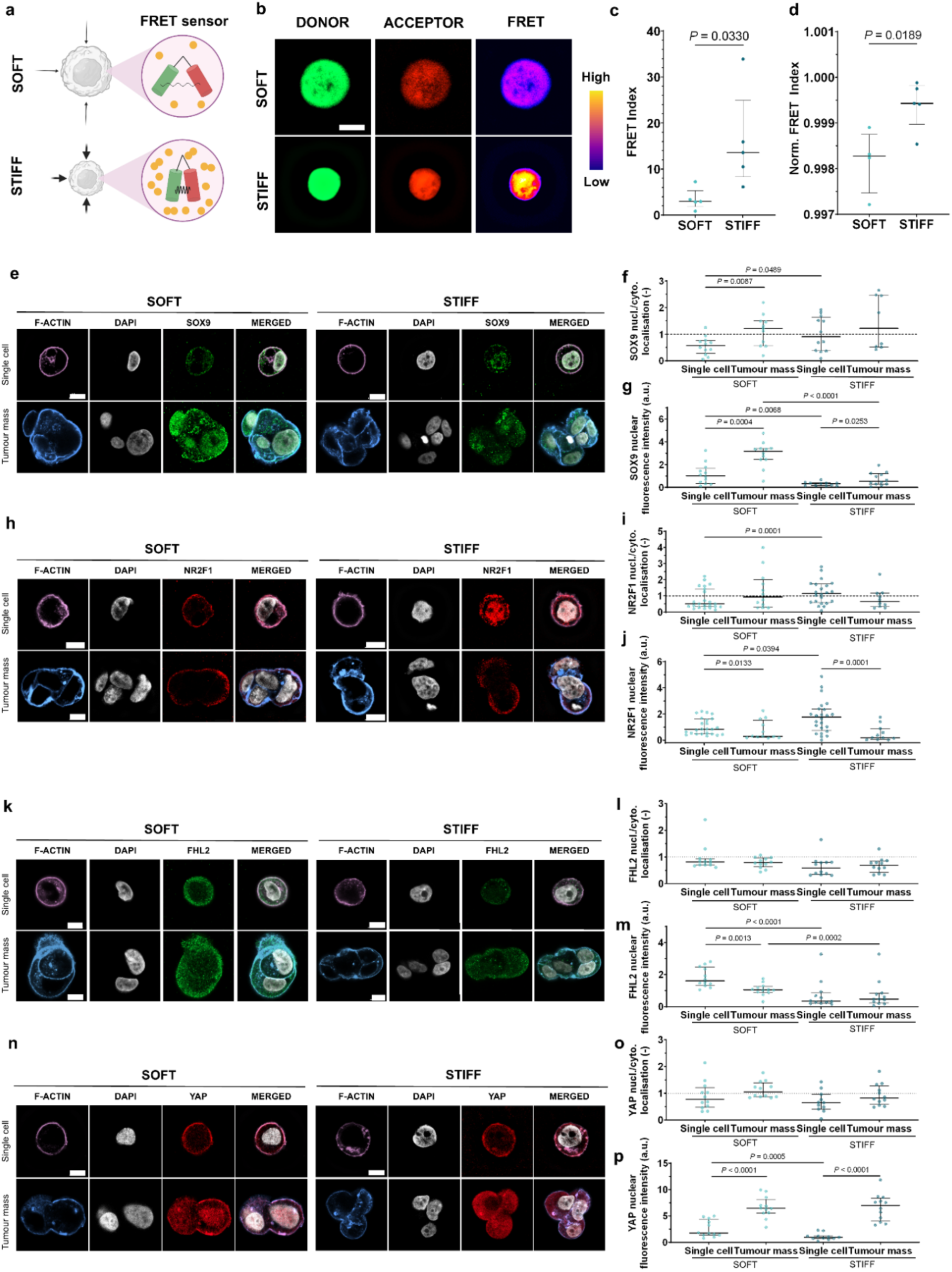
Macromolecular crowding and dormancy-associated and mechanosensitive transcription factors in MCF-7 cells in soft and stiff alginate. **a**, Graphical illustration of CRONOS FRET sensor, mNG donor and mScal acceptor poles connected by a flexible linker, illustrated sensing soft and stiff microenvironments. Schematics created with BioRender. **b**, Representative confocal 2D images of MCF-7 cells transfected with CRONOS sensor within soft and stiff alginate at day 5 (green = donor, red = acceptor, heatmap = FRET signal). **c**, Quantification of raw FRET index measured in Fiji and **d**, normalised against the donor mNG fluorescence intensity (n = 5 cells from n = 3 hydrogels per group). **e-p**, Representative confocal 2D images of MCF-7 cells classified in single cells and tumour masses, nucleo-cytoplasmic ratio and nuclear intensity quantification (n = 12 - 24 units from n = 3 hydrogels per group), for (**e-g**) SOX9, (**h-j**) NR2F1, (**k-m**) FHL2 and (**n-p**) YAP. Scale bars, 10 μm. Lines and error bars show median and interquartile range, respectively. Statistical significance (two-tailed Student’s t-test (parametric) and Mann-Whitney U-test (non-parametric)). a.u. stands for arbitrary units.

We investigated whether these stiffness-dependent changes in protein macromolecular crowding correlated with the commonly studied SOX9 and NR2F1 transcription factors. Nuclear localisation of SOX9 was predominantly observed in tumour masses under both soft and stiff alginate (Fig. 6e,f), with significantly higher nuclear fluorescence intensity in tumour masses within soft alginate (Fig. 6g), linked by others to stem-like dormant states ^56^. In contrast, only single cells within stiff alginate exhibited nuclear localisation of NR2F1 (Fig. 6h,i), with significantly higher nuclear fluorescence intensity compared to tumour masses in stiff alginate, as well as single cells in soft alginate (Fig. 6j), consistent with a prolonged dormant state reported by others ^17,56^.

We examined FHL2 and YAP mechanosensitive transcriptional regulators. FHL2 showed no detectable nuclear localisation (Fig. 6k-m). Similarly, no significant differences in YAP nuclear localisation were detected; however, significant differences in nuclear intensity values were observed in tumour masses compared to single cells in both conditions (Fig. 6n-p).

### Matrix-induced dormancy persists after retrieval from soft alginate

We retrieved cells from soft and stiff alginate after 5 days, cultured them on 2D TCP for 7 days (day 12 of experiment) and assessed dormancy persistence by RNA-seq and tumorigenic potential in CAM models after additional 7 days (day 19 of experiment) (Fig. 7a).

**Fig. 7.**
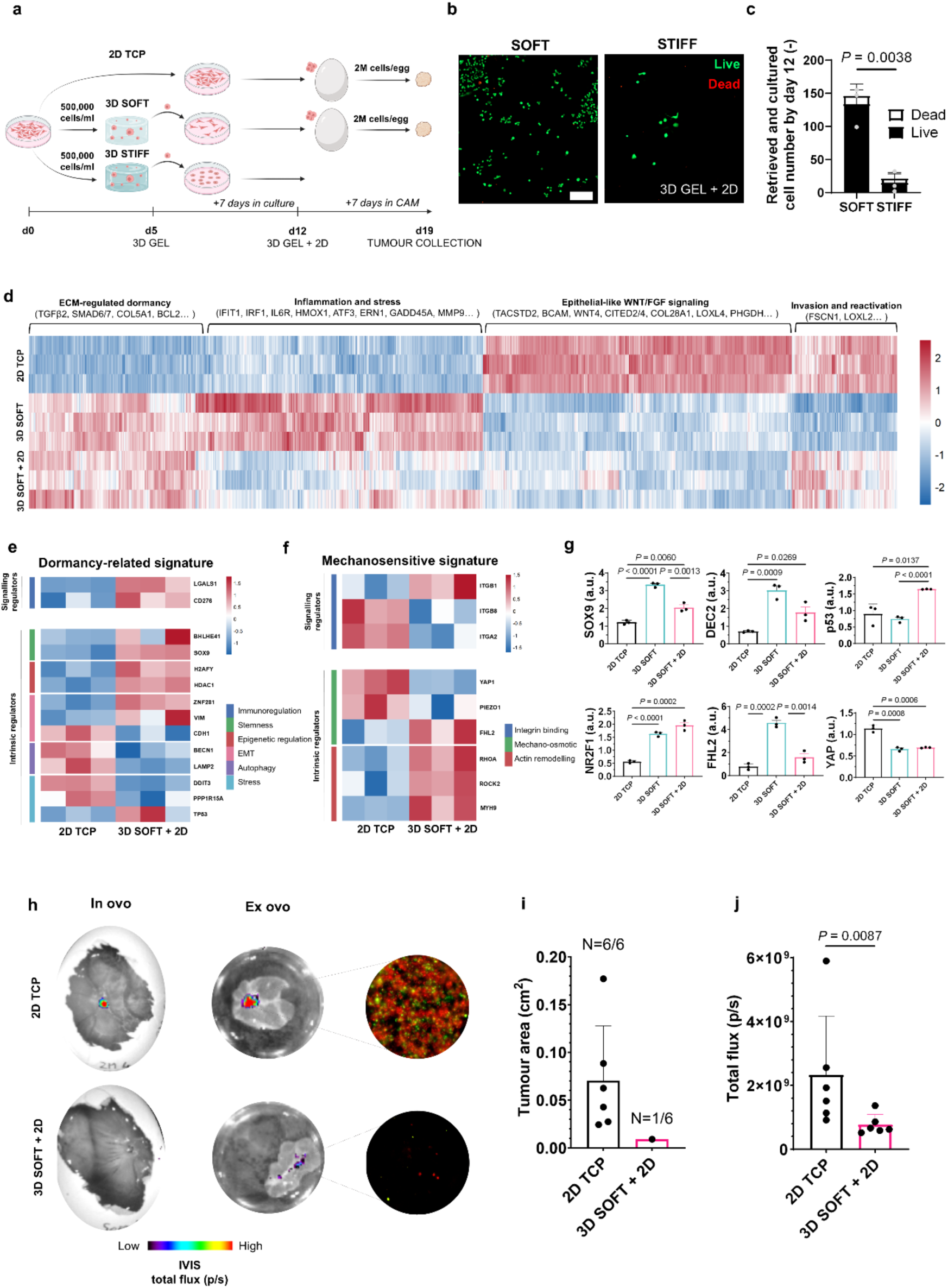
Transcriptomic changes and tumorigenic capacity of MCF-7 cells after exposure to soft alginate. **a**, Graphical illustration of experimental workflow. Schematics created with BioRender. **b**, Representative live/dead (live = calcein, dead = ethidium homodimer) staining of MCF-7 cells on 2D TCP at day 12 after retrieving from soft alginate (10x objective, n = 3). Scale bar, 200 μm. **c**, Quantification of live and dead cell number at day 12 after soft alginate retrieval. **d**, Heatmap of the top 1500 DEGs ranked by absolute log2 fold change (VST) **e-f**, Expression heatmaps of selected candidate genes associated to (**e**) potential dormancy biomarkers and (**f**) mechanosensitive biomarkers, classified into signalling and intrinsic regulators. **g**, RT-qPCR validation of selected transcripts (n=3 biological replicates). **h**, IVIS images of MCF-7 FUCCI2 cells (G_0_/G_1_ = mCherry-hCdt1 and S/G_2_/M = mVenus-hGeminin) CAM assay at day 19, 7 days after cell injection. Fluorescence microscope images of tumour nodules on top of a cover slide (n = 6 eggs per group). **i**, Tumour nodule area quantification in IVIS images from ex ovo images. **j**, Total mCherry fluorescence flux quantification in tumour nodule area. Bars and error bars show mean ± s.d., respectively. Statistical significance (two-tailed Student’s t-test (parametric)). a.u. stands for arbitrary units.

Retrieved cells from soft alginate remained viable (Fig. 7b,c), consistent with a reversible transient dormancy, whereas only an average of 20 cells survived retrieval from stiff alginate (Fig. 7c), despite the small decline in viability during encapsulation through 14 days (Supplementary Fig. 2c).

RNA-seq analyses identified shared matrix-induced dormancy-associated transcriptional signatures in cells retrieved from soft alginate at day 5 (3D soft) and cultured for additional 7 days (3D soft + 2D) (Supplementary Fig. 7a-f, Fig. 7d), while retrieved cells (3D soft + 2D) also upregulated and shared with 2D TCP controls genes associated with invasion and reactivation. GO analysis highlighted protein localisation, autophagy, nucleo-cytoplasmic transport and intrinsic apoptotic signalling as the most enriched biological processes, whereas distinct signatures between both groups confirmed cellular plasticity of retrieved cells (3D soft + 2D) (Supplementary Fig. 7g-i). Retrieved cells (3D soft + 2D) also retained genes associated with immunosuppression (*Lgals1, Cd276*), dormancy-associated stemness and plasticity (*Bhlhe41, Sox9*), EMT (*Vim, Znf281*) and tumour suppression (*Tp53*) (Fig. 7e).

Mechanosensitive genes revealed downregulation of *Itgb8* and *Itga2,* sustained *Itgb1* expression, reduced *Yap1* and *Piezo1* expression, and increased actin cytoskeletal contractility genes (*Rhoa, Rock2, Myh9*) (Fig. 7f). RT-qPCR confirmed persistent upregulation of dec2 (*Bhlhe41*), *Sox9*, *Tp53* and *Nr2f1* (Fig. 7g).

To determine whether matrix-induced dormancy affected tumorigenic potential, retrieved MCF-7-FUCCI2 cells were injected into CAM models (2 · 10^6^ cells/egg) (Supplementary Fig. 8a,b). Only one of six embryos (3D soft + 2D) generated a detectable small tumour nodule with significantly lower fluorescence intensity than 2D TCP control tumours, confirmed by *in ovo* and *ex ovo* IVIS imaging (Fig. 7h-j, Supplementary Fig. 8c,d). These findings demonstrate the persistence of matrix-induced dormancy in transient dormant cells, thereby limiting tumour formation.

### Soft matrices induce therapy resistance in dormant patient-derived cells

Having established that soft alginate induces stem-like transient dormant phenotypes in MCF-7 cells, we assessed the cytotoxic effect of tamoxifen, a selective oestrogen receptor modulator, recapitulating dormant versus proliferative phenotypes in patient-derived samples through controlled biophysical cues.

Primary cells from luminal BC patients (n = 4) were collected from Breast Radiology Services at Hospital Universitario Donostia (HU Donostia) and Basurto (HU Basurto), with different HER2 status according to pathological data (Supplementary Table 1). Score 1+ tumours are classified as HER2-negative (HER2^-^). Score 2+ tumours show equivocal HER2 expression and require in situ hybridisation (ISH) to determine whether they are HER2-positive (HER2^+^). Score 3+ tumours are classified as HER2^+^.

We first enriched cells for stemness to select pan cytokeratin-positive (PanCK^+^) cancer cells and stimulated them with β-estradiol before encapsulation (Fig. 8a). We validated SOX9 expression after 5 days of encapsulation prior to treatment, and confirmed significantly increased nuclear expression in dormant cells within soft alginate compared to proliferative cells in soft GelMA (Fig. 8b,c). We then treated samples with tamoxifen (10 and 50 µM) for additional 2 days.

**Fig. 8.**
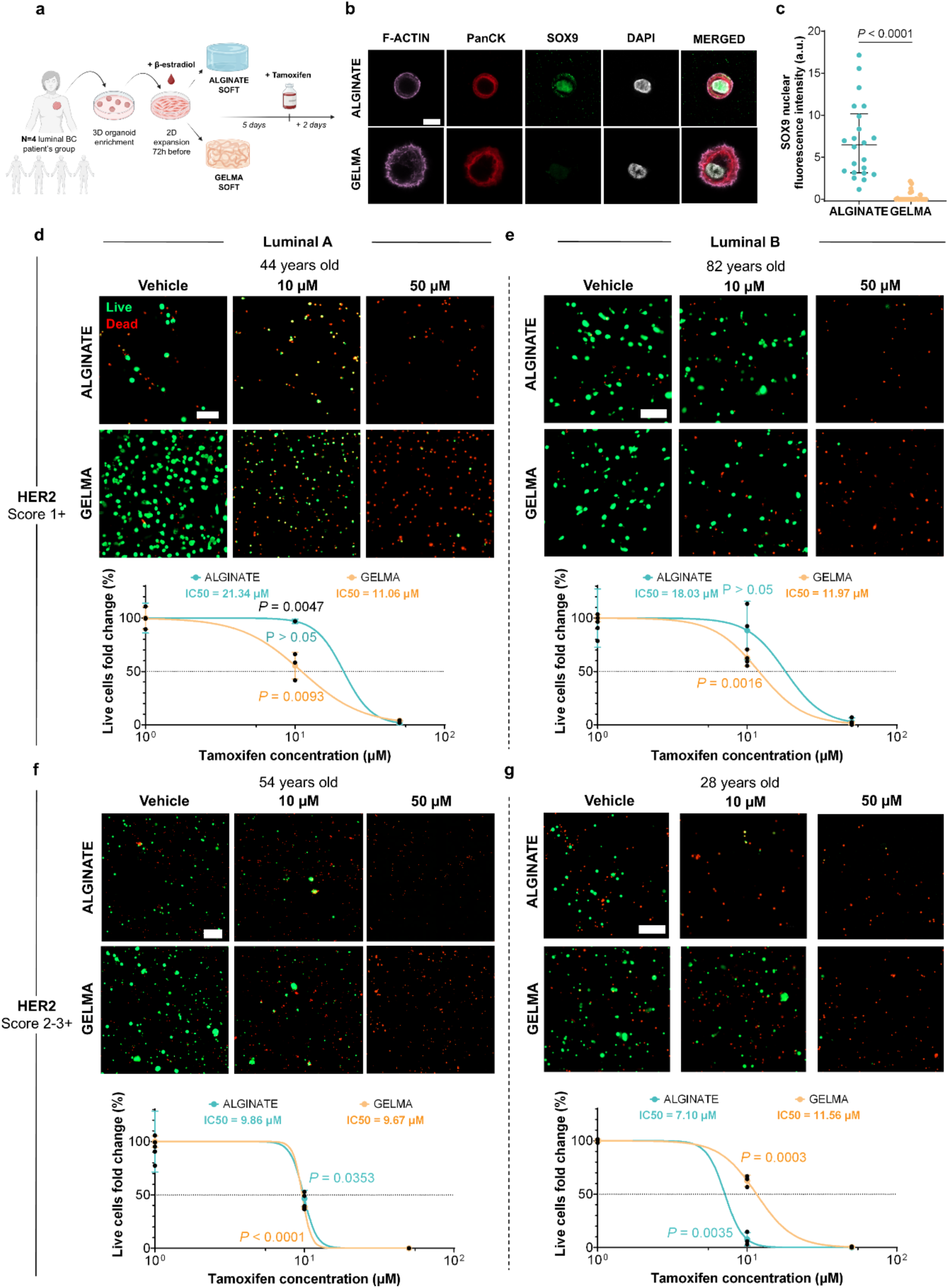
Proof-of-concept of drug testing with patient-derived luminal BC cells. **a**, Graphical illustration of experimental workflow. Schematics created with BioRender. **b**, Representative confocal 2D images of SOX9 expression in PanCK^+^ single cancer cells within 3D soft alginate and GelMA. **c**, Nuclear SOX9 intensity quantification (n = 22-28 nuclei, n = 2 donors, median and interquartile range). **d**, Representative live/dead (live = calcein, dead = ethidium homodimer) maximum projection fluorescence images of cells derived from a group of luminal BC patients (n = 4), classified as luminal A and B, and encapsulated within soft alginate and GelMA, for 5 days followed by 2-day tamoxifen treatment (10 and 50 µM) (n = 3 hydrogel per group). Scale bar, 200 µm. Normalised live cell fold change relative to vehicle controls and corresponding IC50 values derived from non-linear regression curves. Bars and error bars show mean ± s.d., respectively. Statistical significance (two-tailed Student’s t-test (parametric) and Mann-Whitney U-test (non-parametric)).

Drug-induced cytotoxicity was assessed by viability readouts normalised to vehicle controls. In HER2 score 1+ samples, dormant cells showed no difference in the number of live cells when exposed to 10 µM compared to vehicle control, whereas proliferative cells showed a significant viability loss (Fig. 8d). This effect was partially attenuated in the luminal B (more aggressive) donor compared to luminal A patient. At 50 µM, cells from both donors were dead. Consequently, half of the maximum inhibition concentration of the drug (IC50) was higher for dormant cells (19.69 ± 2.34 µM) compared to proliferative cells (11.52 ± 0.64 µM) (Fig. 8d,e).

In HER2 score 2-3+ samples, this drug resistance effect disappeared for the luminal A patient, while the luminal B donor showed a reversal, with greater sensitivity of dormant cells (8.48 ± 1.95 µM) than proliferative cells (10.62 ± 1.34 µM) (Fig. 8f,g).

Overall, soft alginate matrices recapitulate transient dormant states in patient-derived cells and demonstrate matrix-induced therapeutic responses, with SOX9^+^ transient dormant cells, with HER2^low^ expression, exhibiting increased resistance to tamoxifen compared with proliferative cells.

## DISCUSSION

In this study, we engineered tuneable 3D hydrogels that decouple matrix stiffness from adhesion and degradability to investigate how ECM biophysical cues regulate luminal BC dormancy. This approach revealed that soft alginate promotes partially reversible stem-like transient dormancy associated with endocrine therapy resistance, validated through direct drug testing in patient-derived cells. In contrast, stiff alginate promotes autophagic and stress-associated prolonged dormancy, characterised by macromolecular crowding of proteins (Fig. 9). These findings establish matrix biophysical cues as regulator of distinct dormant states, linking ECM mechanics to dormancy regulation and therapeutic response.

**Fig. 9.**
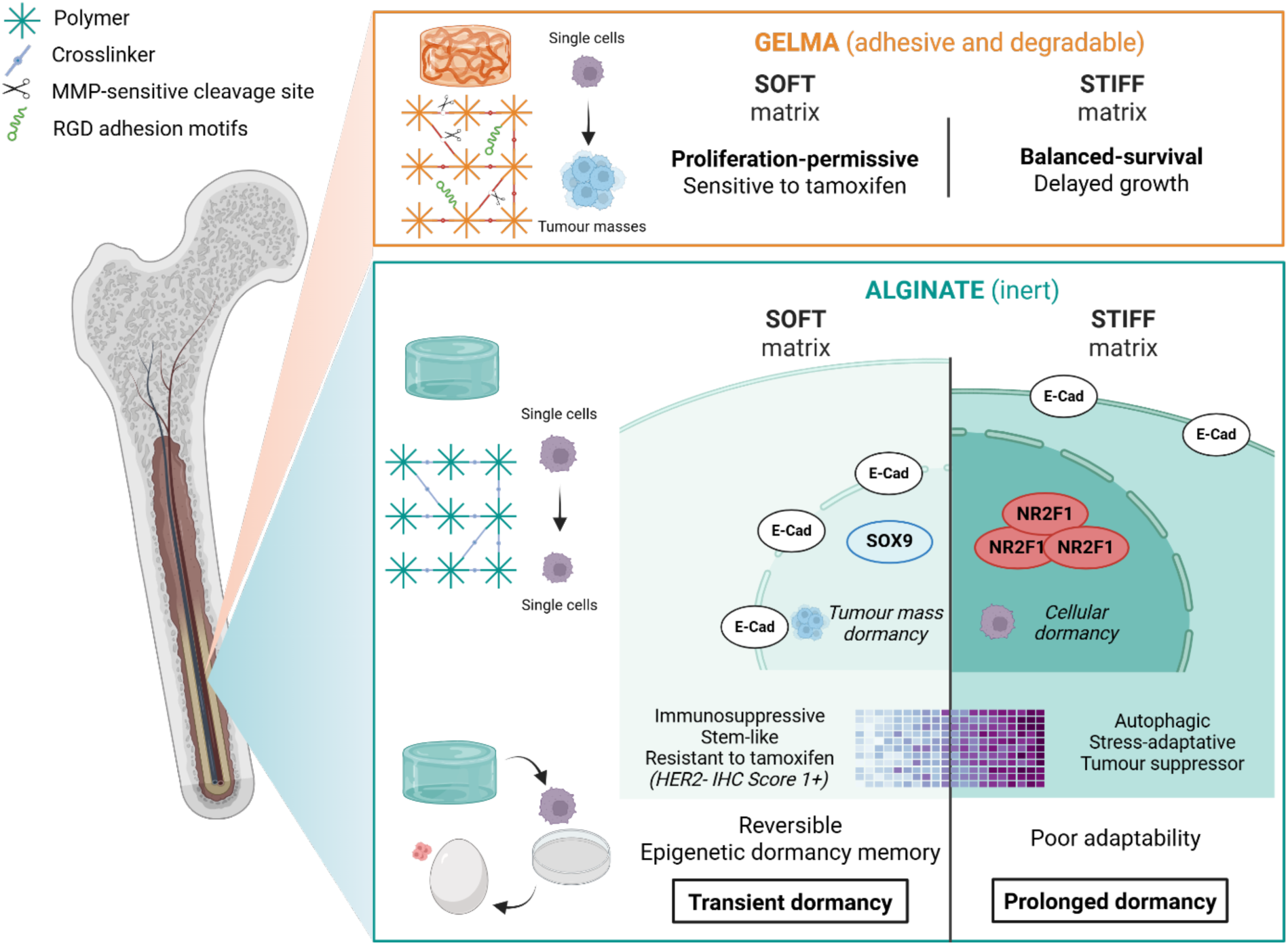
Graphical abstract. A working model illustrating how tuneable 3D bioengineered matrices use different biomaterials to decouple matrix stiffness from adhesion and degradability, thereby regulating transient and prolonged dormant states in the bone marrow, while enabling therapeutic response testing in patient-derived dormant cells.

Using stiffness-matched elastic hydrogels as reductionist models, we isolated the effects of adhesion/degradability and stiffness in dormancy regulation. In soft matrices (1 kPa), GelMA promoted proliferative and metabolically active rounded epithelial-like tumour masses, whereas non-adhesive and non-degradable soft alginate favoured growth-arrested single cells with low metabolic activity and elongated morphologies, with perinuclear E-cadherin localisation, features associated with dormancy ^29,48,50^. Increasing the stiffness, these differences were attenuated (10 kPa), with stiff GelMA delaying tumour growth ^57^ and yielding a balanced distribution of single cells and tumour masses, while stiff alginate induced early volume reductions, consistent with mechanical constraints in inert microenvironments.

Thus, volumetric and morphological features support dormancy-associated phenotypes, together with Ki67/p27^+^ phenotypes in alginate, independent of immunogenic and angiogenic signalling and driven solely by restrictive matrix conditions. Matrix stiffness further modulated stress-associated dormancy in luminal MCF-7 cells, reflected by lower P-ERK:P-p38 ratios in soft matrices. However, this ratio alone was insufficient to characterise dormancy, as it was similarly decreased in soft GelMA, underscoring the need for combination with Ki67^-^/p27^+^ markers.

We intentionally designed non-adhesive alginate hydrogels to induce dormancy, consistent with *in vivo* studies showing that loss of β1-integrin or focal adhesion kinase (FAK) signalling promotes dormancy ^22,58^. Beyond regulating dormancy fate, inert alginate uncovered stiffness-dependent atypical intracellular responses, with stiff alginate inducing reduced cell volume, enlarged nuclei and increased macromolecular crowding, consistent with mechanical confinement and increased intracellular osmolarity ^59–62^.

Macromolecular crowding effects have also been associated with WNK1 osmostress regulation ^55^. Similarly, soft hydrogels showed progressive water uptake (Supplementary Fig. 9), and upregulated the osmoregulatory regulator *Piezo1*, suggesting passive nucleocytoplasmic transport of proteins driven by osmotic adaptations ^63^.

Transcriptomic analyses further demonstrated that soft alginate promotes responsiveness to microenvironmental changes, characterised by upregulated immunosuppressive ^64^ and glycol checkpoints ^65^, together with actin cytoskeleton remodelling, stemness and EMT genes. In contrast, stiff alginate downregulated these programs while promoting NR2F1^high^-associated prolonged dormancy, characterised by autophagy and stress, together with *Tp53*, p21 (*Cdkn1a*) and *Zfp281* prolonged dormancy genes ^66,67^. Although transcriptional profiles were assessed at day 5, these distinct dormancy programmes were predictive of transient and prolonged dormant phenotypes, supporting ECM stiffness as a key regulator of dormancy fate and expanding the concept of permissive versus restrictive microenvironments to soft (transient) versus stiff (prolonged) dormancy-inducing compartments ^67^. This distinction parallels concepts from evolutionary ecology, where dormancy in animals and plants can occur either as shallow (short-term) or deep (long-term) dormancy states, highlighting analogous adaptative dormancy strategies in nature and cancer ^68^.

YAP signalling was consistently suppressed at both transcript and protein levels in dormant luminal MCF-7 cells, particularly in stiff alginate, in agreement with previous evidence that NR2F1 suppresses *Yap1* activity ^69^. Conversely, although *Fhl2* transcripts were upregulated in soft and stiff alginate, and after retrieval, consistent with improved outcomes for luminal BC subtype (Supplementary Fig. 10a), no corresponding protein activation was detected. This contrasts with TNBC models, in which FHL2 appeared activated as a mechanosensitive marker of dormancy ^29^ and associated with poor relapse-free survival (Supplementary Fig. 10b). Notably, nuclear FHL2 activation in MDA-MB-231 cells occurred independent of *Tp53*, whereas in MCF-7 *Tp53* was downregulated during transient dormancy, suggesting its role as a dormancy fate and biomarker for residual disease in the clinic for luminal BC ^70^. These findings highlight the need to explore alternative mechanosensitive pathways, including osmotic stress regulators such as Piezo1, as potential common mediators of dormancy across BC subtypes.

RNA-seq revealed the enrichment of epigenetic regulatory pathways in soft alginate, suggesting persistent chromatin remodelling. Retrieved cells retained dormancy memory, despite partial reactivation, characterised by adult stem (*Sox9*^high^), tumour suppression (*Tp53*^high^) and dormancy (*Nr2f1*^high^, *Znf281*^high^) signatures, consistent with slow-cycling behaviour and limited tumour formation in CAM models.

Given the established role of SOX9 in tamoxifen resistance in luminal BC ^19^, we assessed endocrine therapy response over a 7-day experiment with patient-derived luminal BC cells. Soft alginate induced SOX9^+^ dormant phenotypes associated with higher IC50, particularly in HER2^low^ samples, indicating enhanced endocrine therapy resistance in dormant patient-derived cells. This effect was diminished in HER2 score 2-3+ samples, highlighting subtype-dependent therapeutic resistance. These findings suggest that hydrogel stiffness should be tailored to different BC subtypes, to model heterogeneous dormant states and personalised therapeutic strategies.

To conclude, we established tuneable 3D-bioengineered matrices that disentangle the contributions of ECM biophysical cues to distinct dormancy states, providing mechanistic insights into matrix-induced dormancy and therapeutic responses in short-term drug screening in patient-derived cells.

## METHODS

Materials and reagents (with their respective acronyms) used in this study are summarised in Supplementary Table 2. Theoretical formulas are collected in Supplementary Table 3.

### Alginate functionalisation by carbodiimide chemistry

Norbornene-modified alginate was synthesised as previously described ^71–74^. 5-norbornene-2-methylamine was added at a theoretical DS_theo_ of > 500 molecules per alginate chain.

### Proton nuclear magnetic resonance (^1^H NMR) characterisation

DS_real_ was assessed by NMR measurements performed using an Ultra Shield Plus 500 MHz spectrometer (Bruker; 64 scans), with samples dissolved at a final concentration of 1.5 % (w/v) in D_2_O ^75^. We used unmodified gelatine for GelMA control. Spectra were analysed using MestreNova software (v12.03).

### Casting of photocrosslinkable GelMA and alginate-based hydrogels

Norbornene-modified alginate hydrogels were prepared by a photoinitiated thiol–ene reaction using DTT as a dithiol crosslinker. Alginate was dissolved in 1X PBS overnight at RT under shaking, then mixed with LAP photoinitiator at final concentrations of 2 % and 0.15 % (w/v), respectively. Mechanical properties were tuned by varying DTT concentration: 0.02 mg/mL and 0.25 mg/mL for soft and stiff groups, respectively.

LAP and DTT stock solutions were dissolved in PBS prior to mixing. GelMA hydrogels were fabricated from a commercial gelatin methacryloyl. GelMA was dissolved in PBS for 15 min at 50 °C under shaking and mixed with LAP at 0.15 % (w/v). Hydrogel stiffness was adjusted using 5 % and 9 % (w/v) GelMA for soft and stiff groups, respectively.

For 3D cell encapsulation, cell suspension (5 · 10^5^ cells/(mL gel)) in media (described below) was mixed into the precursor solutions and gently mixed before UV exposure. Solutions were pipetted with positive-displacement pipettes into 1 - 2 mm height custom-built silicone chambers on top of a borosilicate glass plate and covered immediately with a previously treated glass slide to prevent gel adhesion. We used a diameter of 4 mm for regular experiments, 5 mm for immunofluorescence assays and 8 mm for rheology.

The mixture was exposed to UV light for 1 min at 10 mW/cm^2^ (365 nm; OmniCure S2000). Gels were then plated in culture media in 24-well plates until further use. Alginate hydrogels were washed with 1X PBS to remove residual DTT before culture.

### Mechanical characterisation

Rheology measurements were performed to calculate the storage (G’), loss (G’’) and complex (G*) moduli of the gels with and without (replaced with media) cell encapsulation. Frequency sweep measurements were performed from 0.03 to 3 Hz at 3 % shear strain (under linear viscoelastic conditions) with an 8 mm parallel plate geometry using a rheometer (Physica MCR 101; Anton Paar), while keeping the temperature at 37 °C with a Peltier temperature module. Once contact with the gel surface was established, a pre-compression of 10 % of the height of the hydrogel was applied prior to the measurement. No additional hydration was needed as the experiment conditions lasted less than 10 min. Prior to frequency sweep measurements, the elastic range of the hydrogels was assessed by a deformation sweep from 0.001 to 100 % strain at 1 Hz. E modulus was then calculated using Rubber’s elasticity theory in Supplementary Table 3. The mesh size (*ξ*) of hydrogels was approximated based on the average storage modulus G’ at low frequencies (0.03-1Hz).

### Volumetric and mass swelling characterisation

Hydrogel volume changes, wet weight changes and swelling ratios were tracked over time at day 1 (after equilibration), 5 and 14 days, as previously described ^71,73,74^. Hydrogel height and diameter were measured with a calliper, and the volume was calculated assuming a cylindrical shape of hydrogels. For wet weight measurements, hydrogels were weighed at each time point, then freeze-dried and weighed again to obtain the dry weight.

### Cell culture

MCF-7 (HTB-22; ATCC) human BC cell line for the luminal subtype was cultured in low glucose DMEM with 10 % (v/v) FBS, 1 % (v/v) P/S and 1 % (v/v) L-Glutamine. Cells were incubated in a 5 % CO_2_ environment at 37 °C and passaged every 3 - 5 days. Cells were tested for mycoplasma by PCR-based mycoplasma detection assay every month and consistently confirmed negative. Same conditions were preserved for MCF-7 FUCCI2 cells, with lentiviral particle production, transduction and generation previously reported ^29,32^.

Prior to 3D encapsulation, cells were exposed to 0.05 % trypsin-EDTA at 37 °C for a maximum of 5 min for cell detachment. After centrifugation, MCF-7 cells were filtered through a 40 µm cell strainer to obtain a single-cell suspension.

### Cell viability, metabolic activity and morphometric staining

Live and dead cells were stained with 16 µM of calcein AM and 4 µM of EthD-1 dissolved in PBS, followed by a 15 min incubation at 37 °C in the dark. Metabolic activity was assessed using 3D CellTiter-Glo.

Hydrogels were transferred to 24-well plates containing lysis solution and incubated following manufacturer’s instructions. Then, 100 µL of the supernatant was transferred to white 96-well plates and bioluminescence was measured using a GloMax Discover plate reader (Promega).

Cell morphology was assessed with nuclei and actin cytoskeleton using DAPI and Phalloidin, respectively. All steps were performed under orbital shaking in a 24-well plate and using a volume of 500 µL per gel. Gels were washed with PBS and fixed with 4 % formaldehyde for 30 min at RT, followed by a wash with 5 % (w/v) BSA in PBS and permeabilisation with 0.1 % (v/v) Triton X 100 in PBS for 15 min, with a final washing (2X) step with 5 % BSA in PBS. Phalloidin Atto-633 at 0.2 μM concentration was incubated for 2 h at RT protected from light diluted in a 3 % BSA + 0.1 % Triton X 100 buffer solution under mild shaking. A washing step was performed twice with 5 % BSA in PBS before DAPI incubation at 1 μg/mL concentration for additional 30 min with the same buffer and conditions. Finally, samples were washed with PBS (3X) prior to imaging.

### Immunofluorescence staining

All steps were performed under orbital shaking in a 24-well plate and using a volume of 500 µL per gel. Gels were washed with PBS and fixed with 4 % formaldehyde for 30 min at RT. Gels with 5 x 2 mm dimension were then cut in half to facilitate antibody penetration to cells in the centre of the gel. Next, half of the gels were washed with 3 % BSA and permeabilised with 0.3 % Triton X 100 for 15 min with a final washing (2X) step with 3 % BSA in PBS. Primary antibody staining was performed at 4 °C overnight, diluted in a 1 % BSA + 0.3 % Triton X 100 buffer solution. Gels were then washed (3X) with 3 % BSA in PBS for 30 min and incubated with secondary antibody in combination with the corresponding Phalloidin reagent to prevent fluorescence crosstalk between detection channels: 0.2 μM Atto-488, Atto-550 or Atto-633 Phalloidin at RT for 2 h with the same dilution buffer. Followed by washing (3X) with 3 % BSA, DAPI was incubated at RT for 30 min. Samples were washed with PBS (3X) prior to imaging. The primary and secondary antibodies used are listed in Supplementary Table 4.

### Cell retrieval, RNA extraction and purification

Alginate hydrogels were incubated with 1 mg/mL alginate lyase in 10X HBSS for 30 min at 37 °C under mild rocking, pipetting gels up and down every 10 min. Pellet was then collected, centrifuged and washed (2X) with HBSS and cultured on 2D TCP with growth media for the following experiments.

For RNA extraction and purification, we used Maxwell RSC simplyRNA Cells Kit following manufacturer’s instruction. Briefly, cell pellet was lysed using 200 µL of 1-Thioglycerol/Homogenisation solution. Before loading samples into the cartridges, 200 µL of lysis buffer was added to lysed cells and vortexed for 15 seconds. 400 µL of lysate were transferred to the cartridge followed by 5 µl of DNase to the corresponding column. After one hour of RNA extraction and purification in Maxwell RSC Instrument (Promega), samples were collected and stored at −80 °C until further use.

### RNA sequencing and analysis

The quality and quantity of the RNA were assessed using Qubit RNA HS Assay Kit and Agilent RNA 6000 Nano Chips, respectively. Strand-specific mRNA libraries were prepared using the Illumina Stranded mRNA Prep kit, according to manufacturer’s instruction. Libraries were quantified using Qubit dsDNA HS Assay Kit and quality-checked on Agilent 2100 Bioanalyzer using Agilent High Sensitivity DNA Kit. Libraries were sequenced in paired-end mode at the Genome Analysis Model of CIC bioGUNE (Derio, Spain) under the supervision of Ana M. Aransay, generating approximately 40 million 2 x 150 bp reads per library.

Transcript abundance was quantified using Salmon (v1.10.2) ^76^, with the human *Ensembl GRCh38* cDNA transcriptome reference (Ensembl release 86). Transcript-level estimates were summarised to gene-level counts with tximport (v1.38.2) ^77^ in R (v4.5.3). DEGs analysis was performed using DESeq2 (v1.50.2) ^78^. Raw counts were normalised using the median-of-ratios method, dispersion parameters were estimated under a negative binomial model, and statistical significance was assessed using Wald tests followed by Benjamini– Hochberg (NH) correction for multiple testing. Log_2_ fold changes were shrunk using *apeglm* ^79^ or *ashr* ^80^ methods. Genes with a BH adjusted p < 0.05 were considered significantly differentially expressed.

Quality control included assessment of library size, variance-stabilising transformation (VST) ^81,82^, principal component analysis (PCA) and hierarchical clustering based on Euclidean distances between samples.

Functional enrichment analysis was performed using Gene Ontology biological process terms using clusterProfiler (v4.18.4) ^83^.

### RT-qPCR

After RNA quantification using an EPOCH spectrophotometer, RNA was converted to cDNA using the Maxima First-Strand cDNA synthesis kit with incorporated DNAse treatment according to the manufacturer’s instructions. qPCR was performed using TB Green Premix Ex Taq with a final forward and reverse primer concentrations of 1 µM and cDNA concentrations of 0.5 ng/µl. qPCR was conducted for up to 40 cycles on a CFX 384 PCR system (Bio-Rad). Gene expression levels were determined using the ΔΔCt method and normalised against three housekeeping genes, GAPDH, CCSER2 and PCBP1 ^84^. Primers were purchased from Condalab and primer sequences used for qPCR are summarised in Supplementary Table 5.

### Molecular crowding sensor (CRONOS) expression

MCF-7 cells were transiently transfected with the pcDNA3.1-CRONOS sensor plasmid ^54^ using Lipofectamine 2000 according to manufacturer’s instructions. In a 6-well plate, 15 µL of transfection reagent and 5 µg/µL of DNA were used per well. All growth media and buffers were prepared without FBS or P/S to improve transfection efficiency. After 48 h, cells were encapsulated in 3D hydrogels and transferred to plates containing growth media with FBS and P/S.

### Tumour growth in chicken chorioallantoic membrane (CAM) models

CAM model assays were carried out according to previously established protocols ^52,75,85,86^. Briefly, fertilised chicken eggs (White Leghorn; Granja Santa Isabel, Spain) were incubated vertically at 37 °C, 55 % humid atmosphere and with 45-minute schedule rotation in Ovation 56 EX Incubator (Brinsea Incubation Specialists, #AG47A). CAM models were opened by puncturing with an electronic diamond scriber the top and side (air bag side close to a venous branching) of the shell and sucking up air from the air bag using a pipette bulb, creating an air pocket on top of the CAM at day 10 of chicken embryo development.

Then, MCF-7 FUCCI2 cells in 2 · 10^6^ cells/50 μl of cell suspension per egg were inoculated with a pipette on top the membrane at day 10 of chicken embryo development. Beforehand, cells were trypsinised and their concentration was determined with Countess II (Invitrogen) using 0.4 % trypan blue stain. Viable cells were prepared in cell suspensions in PBS with 1 % P/S mixed with Matrigel in 1:1 dilution ratio.

Eggs were incubated at 37 °C in a 55 % humid atmosphere for 7 days before tumour collection. At day 17 of chicken embryo development, eggs were opened and tumours were imaged.

### Patient-derived sample collection, enzymatic digestion and 3D encapsulation

Samples and data from patients included in this study were provided by the Basque Biobank www.biobancovasco.bioef.eus (National Registry of Biobanks (B.0000140)), integrated in the Model ISCIII Biomodels and Biobanks (PT23/00013) and processed under ethical approval from the Ethics Committee of the Basque Country (CEIm-E, Code PI+CES-BIOEF 2024-13). Core needle biopsies from luminal BC patients were collected in Breast Radiology Services from HU Donostia and HU Basurto and recovered into a centrifuge tube in MACS Tissue Storage Solution supplemented with 1 % P/S until further use.

Non-tumoral tissue was removed using disposable blades and tumour tissue was further minced to approximately 1 mm^3^ fragments, followed by enzymatic digestion with collagenase/hyaluronidase or collagenase type II digestion at 1:10 dilution or 1 mg/mL in basal media, respectively, for 60 min at 37 °C under shaking conditions. Basal media formulation was Advanced DMEM-F12 with 1 % (v/v) HEPES, 1 % (v/v) P/S and 1 % (v/v) Glutamax.

Samples were then filtered through pre-wetted 100 μm and extra 70 μm cell strainers, centrifuged (5 min at 500 g and 4 °C) and treated for red blood cell lysis with ACK Lysing Buffer at RT for 3 min. Cells were washed (1X) with PBS and cultured in different conditions depending on the sample type (Supplementary Table 6). Cells in 2D were cultured in adherent 6-well plates, while 3D organoids were maintained in non-treated 24-well plates with the corresponding complete media with 2% (v/v) growth factor-reduced Matrigel and cultured under suspension conditions.

For 3D hydrogel encapsulation, cells were first cultured in complete media with 100 nM β-estradiol and then encapsulated as described above.

### Drug testing

To assess patient-derived cells response to endocrine therapy against selective oestrogen receptor modulators, tamoxifen was diluted at 10 and 50 µM in complete media with β-estradiol. After 5 days of 3D hydrogel encapsulation, different drug dilutions were added to the gels for additional 2-day drug treatment. The vehicle was calculated at the highest concentration of DMSO, which contained 0.5 % (v/v) of it, and did not show any negative effect on cell viability.

### Image acquisition

Epifluorescence imaging of 2D TCP and 3D hydrogels in 24-well plates was performed using a Zeiss AxioObserver 7 inverted microscope. 10x/0.3 NA air objective (Zeiss #420341-9911-000) was used for FUCCI2, Live/Dead, and 2D cultures and 20x/0.4 NA air objective (Zeiss #421351-9972-000) for 3D immunofluorescence images within hydrogels mounted on glass coverslips. For 3D stacks, the focal plane was set at middle height of the hydrogel and z-stacks were acquired over 200 – 400 µm with 20 µm interval steps (10x objective) or 10 µm steps (20x objective), collecting phase-contrast and fluorescence channels (Supplementary Table. 7).

Confocal imaging of 3D hydrogels was performed on a Zeiss LSM 900. 20x/0.8 NA air objective (Zeiss #420650-9902) was used for volumetric and morphometric imaging and 40x/1.3 NA oil objective (Zeiss #420462-9900) for immunofluorescence imaging. For 3D imaging, z-stacks spanning 200 µm were acquired at 1 µm steps (20x objective). For immunofluorescence, hydrogels were sectioned and imaged as 2D cut planes to avoid signal overlap (Supplementary Table. 8). CRONOS images were acquired by exciting the donor mNG at 488 nm laser (1 %) and collecting two simultaneous images of the donor mNG (520 - 560 nm, 700 V) and the acceptor mScal (590LP, 800 V). The laser pinhole was kept at 1 airy unit.

IVIS images of CAM models were acquired using Lumina XRMS Series III (PerkinElmer). For FUCCI2, both fluorescence channels mCherry and mVenus were recorded at 1 s of exposure time. Excitation and emission filters of 560 nm and 620 nm were used for mCherry, respectively, and 500 nm and 570 nm for mVenus. For *in ovo* images, the eggshell was partially opened to capture the intact proximal CAM, whereas ex ovo images were acquired by collecting the proximal CAM and placing it in PBS in 6-well plates.

### Image analysis

Unless otherwise stated, all the microscope image analyses were performed using Dragonfly software, Version 2025.1 for [Windows], Comet Technologies Canada Inc., Montreal, Canada; software available at https://dragonfly.comet.tech/.

Segmentation Wizard tool was used for segmentation models ^87^. Training data was generated by manual labelling of 2D frames (5 frames) and used to train a U-Net model (100 epochs, stride ratio 0.25, batch size 256, Adadelta optimisation ^88^, ORSDiceLoss validation). The trained model was applied to the whole dataset.

2D ROIs were obtained using multi-ROI conversion (Supplementary Fig. 3a). For 3D ROIs, noise was smoothed (kernel size 5) and ROIs in contact with the image border were excluded, along with cells < 1000 µm³ and nuclei < 100 µm³. Datasets were then unified by multi-ROI conversion, and single cell or tumour mass 3D volume and sphericity scalar values were quantified (Supplementary Fig. 3b). For confocal 2D analysis, F-actin and DAPI images were segmented using Otsu thresholding ^89^, followed by smoothing (kernel size 5) and cytoplasm manual labelling. Perinuclear regions were defined by dilating nuclear ROIs (kernel size 11) and subtracting the original nuclear ROI. ROIs were unified by multi-ROI conversion and the mean intensity was measured in raw images (Supplementary Fig. 3c).

CRONOS FRET index calculation was performed using the FRET and Colocalization Analyzer plugging in Fiji software using Java 8 (www.imagej.net).

IVIS images were analysed using Living Image v4.8 (PerkinElmer). The same ROI size was used for each replicate, assuming a small background signal in the surrounding area of each tumour nodule.

Tumour signal area and total photon flux per second (p/s) covering the proximal CAM were quantified only for mCherry signal, as mVenus signal overlapped with the white eggshell noise.

### Statistical analysis

Statistical analysis was performed with GraphPad Prism 10 (version 10.6.1 for Windows, GraphPad Software Inc., Boston, Massachusetts USA). Cell number counts and proportions (%) were corrected with S_t_ volumetric swelling factor (cell number · S_t_) (Supplementary Fig. 9). To test for normality, the Shapiro-Wilk test was applied. For statistical comparison between two groups, two-tailed Student’s t-test or Mann-Whitney U-test were performed for normally-distributed or non-parametric groups, respectively. Statistical analyses were performed after excluding outliers identified using the ROUT method (Q = 10 %).

## REPORTING SUMMARY

Further information on research design is available in the Nature Research Reporting Summary linked to this article.

## DATA AVAILABILITY

The main data supporting the results in this study are available within the paper and its Supplementary Information. All data and image analysis models generated in this study are available on Zenodo with the identifier: 10.5281/zenodo.21461446. RNA-seq data in this publication has been deposited in NCBI’s Gene Expression Omnibus (Edgar *et al.*, 2002) and are accessible through GEO Series accession number GSE341386, with the identifier: https://www.ncbi.nlm.nih.gov/geo/query/acc.cgi?acc= GSE341386.

## ACKNOWLEDGEMENTS

The authors thank group alumni **Claudia Garrido**, **Daniela Garske** and **Hubert Taïeb** for their collaboration in establishing protocols, and current group members **Laura Fallert** and **Rocío Cárdenas** for their assistance with NMR. We are also grateful to **Mercedes Fernandez** and **Itziar Insua** for their support with rheology, to the groups of **Paloma Bragado**, **Veronica Torrano**, **Lorea Valcarcel** and **Manuel Salmeron** for their assistance with CAM experiments and protocols, **Ana M. Aransay** from CIC bioGUNE for RNA-seq service and **Pedro Guerrero** from EHU for lyophilisation of alginate samples. We also thank **Kohsuke Kanekura** (Tokyo Medical University) for sharing CRONOS plasmid. We are also grateful to the **MATRIXinCANCER** (PID2021-123013OB-I00) project network, with special thanks to **Amaia de la Maza** as a Clinical Research Coordinator, **Maria Caffarel’s** group, and platforms and data managers at **Biogipuzkoa** and **Biobizkaia**. We want to particularly acknowledge the patients and the **Basque Biobank** www.biobancovasco.bioef.eus integrated in the Model ISCIII Biomodels and Biobanks (PT23/00013) for their collaboration, as well as the **Breast Radiology Service members** from HU Donostia and HU Basurto. Finally, we thank **Marliette R. Matos** for RNA-seq analysis support.

## FUNDING

U.H and J.B. disclose support for the research of this work from the Basque Government Department of Education (grants PRE_2022_1_0006, PRE_2023_2_0096, PRE_2024_2_0005, PRE_2025_2_0097and PRE_2024_2_0062). U.H. discloses support for the research of this work from the European Association for Cancer Research (EACR) (grant UE/2025/EACR/HERAS) and European Molecular Biology Organisation (EMBO) (grant n°11725). G.d.C. and A.C. disclose support for the research of this work from the Spanish Ministry of Science and Innovation (grants PID2024-159104OB-I00 (G.d.C); and grants PID2021-123013OB-I00 and #PID2025-169199OB-I00 (A.C.), funded by MCIN/AEI/10.13039/501100011033/FEDER, UE). O.M.I. discloses support for the research of this work from Gipuzkoako Foru Aldundia-Diputación Foral de Gipuzkoa (grant 2025-CIE4-000015-01) and Fundación Científica Asociación Española Contra el Cáncer (grant POSTD234723MITX). A.C. discloses support for the research of this work from Fundación Científica Asociación Española Contra el Cáncer (LABAE223466CIPI), Programa FORTALECE - IIS Biodonostia (FORT23/00026), European Research Council Consolidator Grant (grant DORMATRIX, 101123883) and IKERBASQUE Basque Foundation for Science. J.M.P.S., I.T., L.I., I.C., R.A., J.A.G., E.A.B., M.F.C., R.M.Z.S., A.L. and A.R. declare no relevant funding.

## AUTHOR CONTRIBUTIONS

- Study design: U.H., O.M.I., A.C.
- Experimental work: U.H., J.M.P.S., J.B., I.T., L.I., I.C., G.d.C., E.A.B., M.F.C., R.M.Z.S., A.L., O.M.I.
- Data analysis: U.H., J.M.P.S., J.B., O.M.I., A.C.
- Visualisation: U.H., J.M.P.S., J.B., O.M.I., A.C.
- Data interpretation: U.H., J.M.P.S., J.B., I.T., L.I., I.C., R.A., G.d.C., J.A.G., E.A.B., M.F.C., R.M.Z.S., A.L., A.R., O.M.I., A.C.
- Project supervision: O.M.I., A.C.
- Drafting manuscript: U.H., O.M.I., A.C.
- Revising manuscript content and approving final version: all authors

## COMPETING INTERESTS

The authors declare no competing interests.

## ADDITIONAL INFORMATION

**Supplementary information.** This manuscript contains supplementary materials available at: (link)

## Notes

### Competing Interest Statement

The authors have declared no competing interest.

