## Supplementary Information for "Matrix biophysical cues drive transient or prolonged breast cancer dormancy programmes"

**SUPPLEMENTARY FIGURES**

**Supplementary Fig. 1 |** Gelatin methacryloyl (GelMA) and norbornene-modified inert alginate hydrogels characterisation.
**Supplementary Fig. 2 |** Cell viability and ATP release over time within 3D hydrogels.
**Supplementary Fig. 3 |** Deep learning-based 3D image analysis workflow in ORS Dragonfly.
**Supplementary Fig. 4 |** Distinct single and tumour mass distribution across all the conditions and mesenchymal versus epithelial characterisation in stiff alginate.
**Supplementary Fig. 5 |** 3D nuclear volumetric analysis in single cells and tumour masses.
**Supplementary Fig. 6 |** RNA-seq analyses in soft and stiff alginate at day 5.
**Supplementary Fig. 7 |** RNA-seq analyses in MCF-7 cells on 2D TCP, 3D soft alginate and retrieved and post-retrieval cells.
**Supplementary Fig. 8 |** Tumour nodule formation in CAM models.
**Supplementary Fig. 9 |** Mass and volumetric swelling characterisation of GelMA and alginate hydrogels over time**.**
**Supplementary Fig. 10 |** Distinct relapse-free survival of patients for different breast cancer subtypes.

**SUPPLEMENTARY TABLES**

**Supplementary Table. 1 |** Pathological data from a luminal BC patient group (n=4).
**Supplementary Table. 2 |** Materials and reagents.
**Supplementary Table. 3 |** Theoretical formulas.
**Supplementary Table. 4 |** Primary and secondary antibodies.
**Supplementary Table. 5 |** Forward and reverse primer sequences of dormancy genes analysed by RT-qPCR.
**Supplementary Table. 6 |** Culture conditions and media compositions of different group of patients.
**Supplementary Table. 7 |** Zeiss AxioObserver 7 epifluorescence image acquisition data for 3x3 binning.
**Supplementary Table. 8 |** Zeiss LSM 900 confocal image acquisition data with 1 airy unit pinhole aperture.

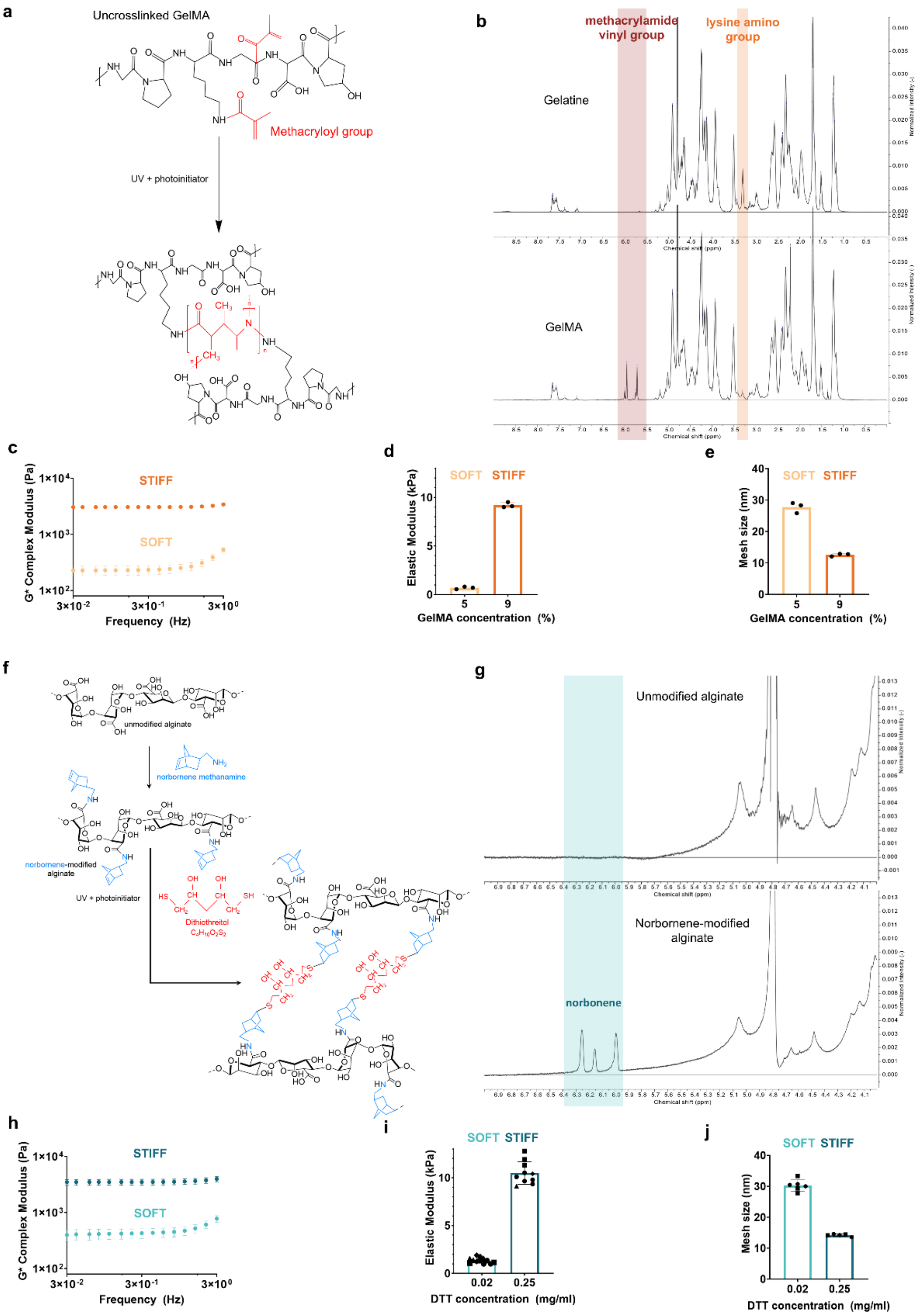


**Supplementary Fig. 1 | Gelatin methacryloyl (GelMA) and norbornene-modified inert alginate hydrogels characterisation. a,** Photocrosslinking of GelMA chains under UV irradiation. Schematics created with ChemDraw. **b,** $^1$H NMR spectra of raw gelatine and GelMA, indicating hydrogens from vinyl and amino groups of methacrylamide and lysine respectively (GelMA source = Porcine Type A, commercial DS = 80 % and $DS_{real}$ = 83 %) **c,** Rheological measurements of GelMA hydrogels showing complex modulus (G*) values for soft (5 %) and stiff (9 %) GelMA hydrogels at day 1 after overnight swelling without cell encapsulation (n=3). **d-e,** Elastic modulus and mesh size derived from average G*. **f,** Norbornene-modified alginate chains crosslinked upon exposure to UV light in the presence of the non-degradable dithiothreitol (DTT) crosslinker by thiol-ene reaction. Schematics created with ChemDraw. **g,** NMR spectra of unmodified and norbornene-modified alginate (molecular weight = 200 kDa, $DS_{theoretical}$ = 500 molecules / alginate, $DS_{real}$ = 107.9 ± 10.42 molecules / alginate, DS (%) = 21.58 ± 2.083 % and reaction efficiency = 21.58 ± 2.083 %; mean ± s.d., n = 3 synthesis). **h,** Rheological measurements of alginate hydrogels showing complex modulus (G*) values for soft (0.02 mg/mL DTT) and stiff (0.25 mg/mL DTT) alginate hydrogels at day 1 after overnight swelling without cell encapsulation (n = 3 synthesis). **i-j,** Elastic modulus and mesh size derived from average G*. Bars and error bars show mean ± s.d., respectively.

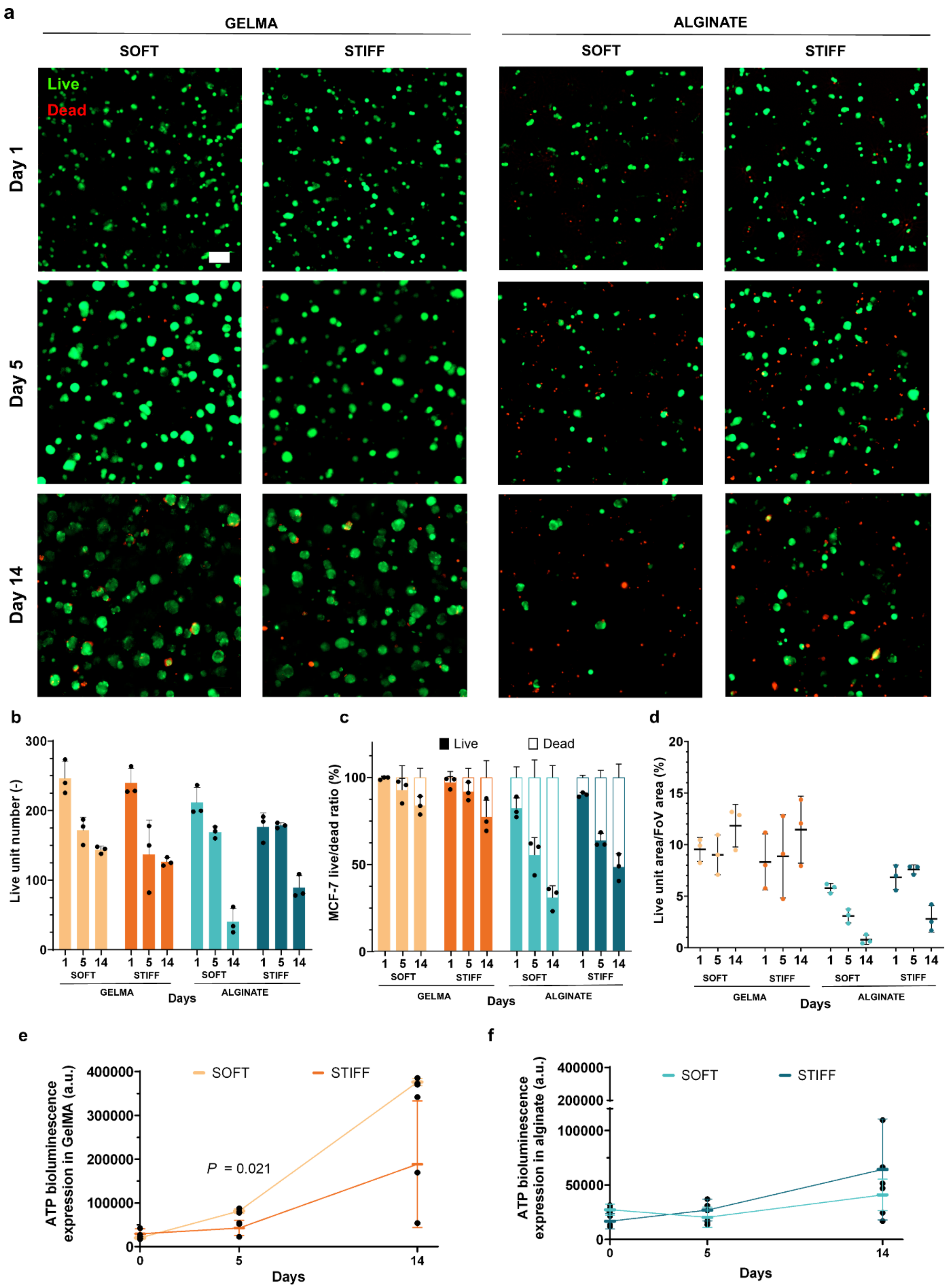


**Supplementary Fig. 2 | Cell viability and ATP release over time within 3D hydrogels. a,** Representative live/dead (live = calcein, dead = ethidium homodimer) maximum projection fluorescence images of MCF-7 cells in soft and stiff GelMA and alginate gels at day 1, 5 and 14. Scale bar, 100 µm. **b,** Live cell units across all groups at day 1, 5 and 14. **C,** Fraction of live and dead cells at day 1, 5 and 14. **D,** Live cell area normalised against field of view area at day 1, 5 and 14. **e-f,** ATP release as a measure of metabolic activity in soft and stiff (e) GelMA and (f) alginate at day 0, 5 and 14 of experiment. Bars and error bars show mean ± s.d., respectively. Statistical significance (two-tailed Student's t-test (parametric). FoV stands for Field of View and a.u. stands for arbitrary units.

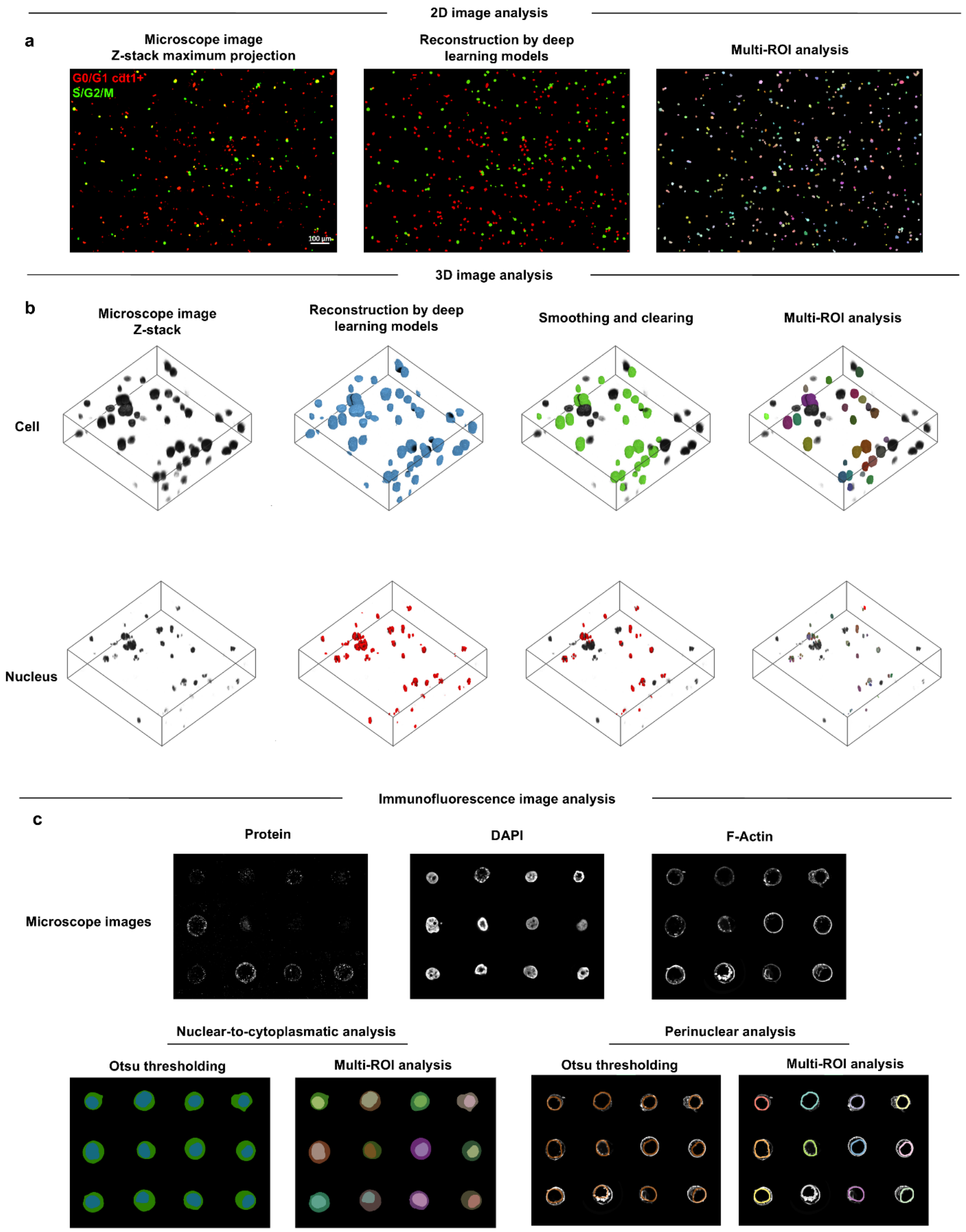


**Supplementary Fig. 3 | Deep learning-based 3D image analysis workflow in ORS Dragonfly**. **a,** 2D analysis for Live/Dead, FUCCI2 or immunofluorescence images using maximum projection z-stacks. Images were reconstructed with custom-made deep learning models and segmented into units using multi-ROI analysis. **b,** 3D volumetric and morphometric analysis from confocal z-stacks of cytoplasm and nuclei reconstructed with deep learning models, followed by smoothing (kernel size 5), clearing multi-ROI segmentation. **c,** 2D mosaic-based immunofluorescence analysis of single cells and tumour masses. Protein, DAPI (nucleus) and F-actin (cytoskeleton) segmentation by Otsu thresholding and multi-ROI analysis, classified between nuclear-to-cytoplasmatic and perinuclear analyses.

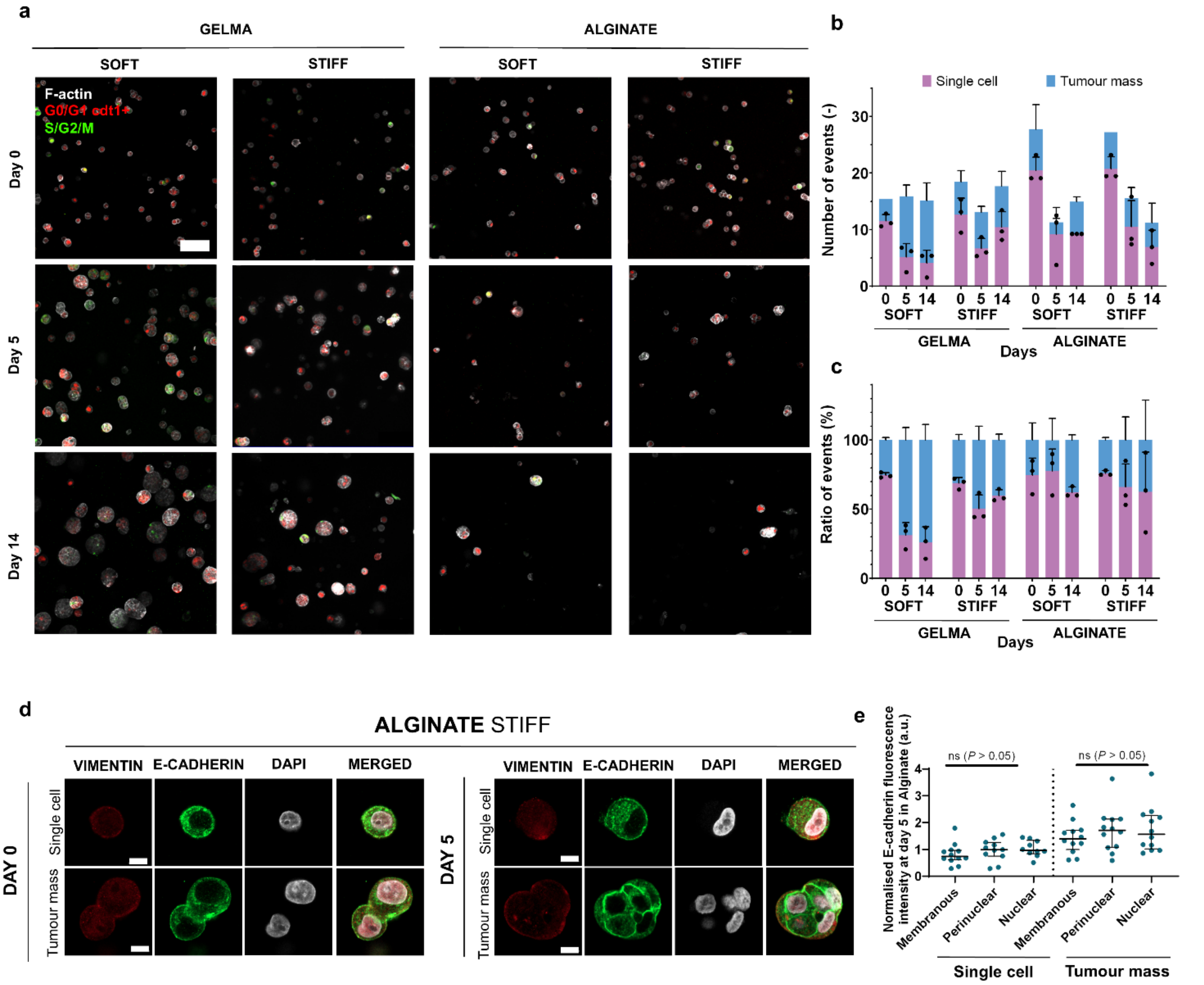


**Supplementary Fig. 4 | Distinct single and tumour mass distribution across all the conditions and mesenchymal versus epithelial characterisation in stiff alginate. a,** Representative confocal MCF-7 FUCCI2 and F-actin staining in GelMA and alginate hydrogels at days 0, 5 and 14 (200 µm z-stacks, 20x objective, n = 3). Scale bar, 100 µm. **b,** Absolute numbers and **c,** fractions of single cells and tumour mass over time in each hydrogel condition (n = 3). Bars and error bars show mean ± s.d., respectively. **d,** Representative confocal 2D images of MCF-7 cells stained for Vimentin (red) and E-cadherin (green) at day 0 and 5 in stiff alginate hydrogels. Cells are classified as single cells and tumour masses (DAPI = white). Scale bars, 10 µm. **e,** Quantification of E-cadherin intensity in membranous, perinuclear and nuclear regions at day 5, normalised to day 0 in stiff alginate (n = 12 units from n = 3 hydrogels per group). Lines and error bars show median and interquartile range, respectively. Statistical significance (two-tailed Student's t-test (parametric) and Mann-Whitney U-test (non-parametric)). a.u. stands for arbitrary units.

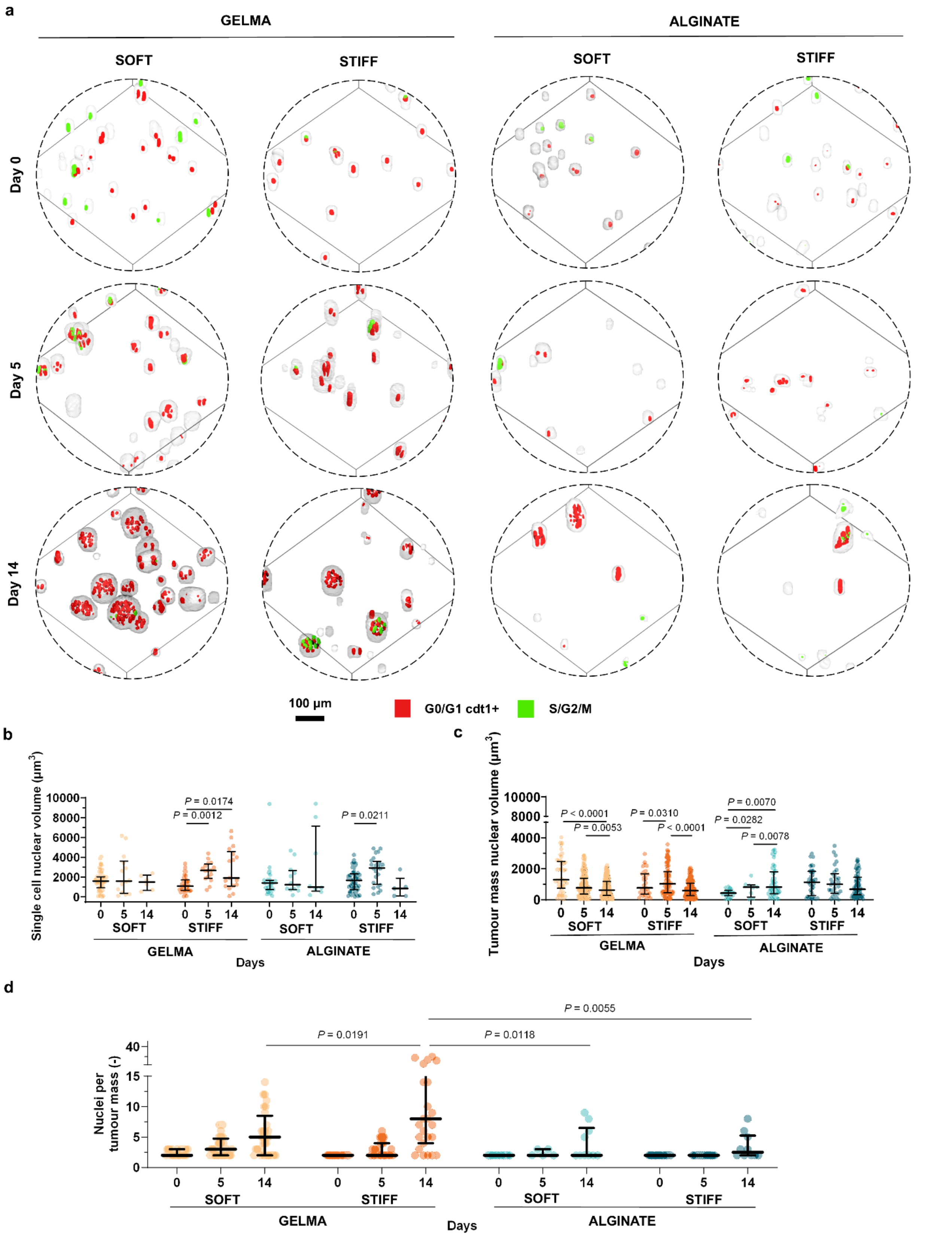


**Supplementary Fig. 5 | 3D nuclear volumetric analysis in single cells and tumour masses. a,** 3D reconstruction of confocal MCF-7 FUCCI2 cells ($G_0/G_1$ cdt1+ = mCherry-hCdt1 and $S/G_2/M$ = mVenus-hGeminin) with F-actin staining (grey) in soft and stiff GelMA and alginate hydrogels at day 0, 5 and 14 (200 µm z-stacks, 20x objective, n = 14 - 392 nuclei from n = 3 hydrogels per group). 3D reconstruction in ORS Dragonfly. **b,** Single cells and **c,** tumour mass nuclear volume quantification at days 0, 5 and 14. **d,** Number of nuclei per tumour mass at days 0, 5 and 14 (n = 5 - 52 clusters from n = 3 hydrogels per group). Lines and error bars show median and interquartile range, respectively. Statistical significance (two-tailed Student's t-test (parametric) and Mann-Whitney U-test (non-parametric)). a.u. stands for arbitrary units.

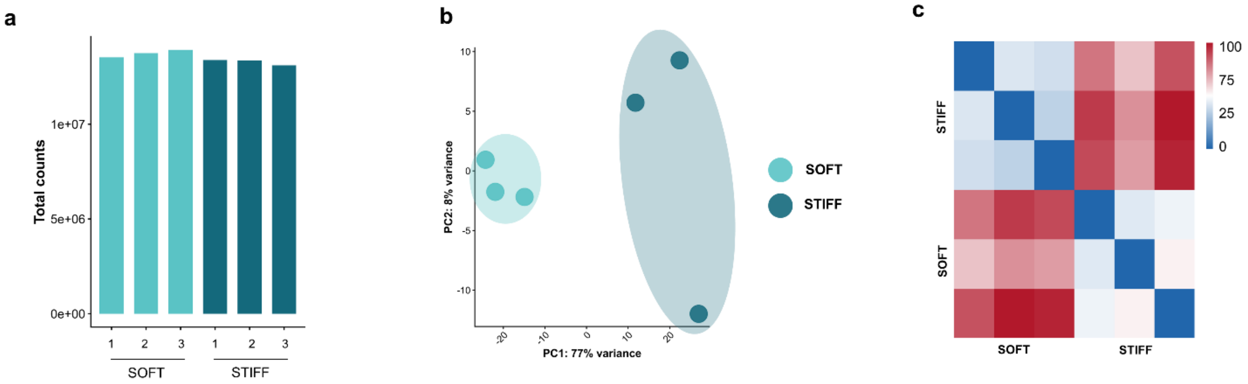


**Supplementary Fig. 6 | RNA-seq analyses in soft and stiff alginate at day 5. a,** RNA-seq library size per samples showing total read counts in soft and stiff alginate **b,** Principal-component analysis (PCA) of variance-stabilised transformed (VST) gene expression profiles between soft and stiff alginate **c,** Sample-to-sample distance matrix based VST normalised counts, showing global transcriptional similarity and clustering structure.

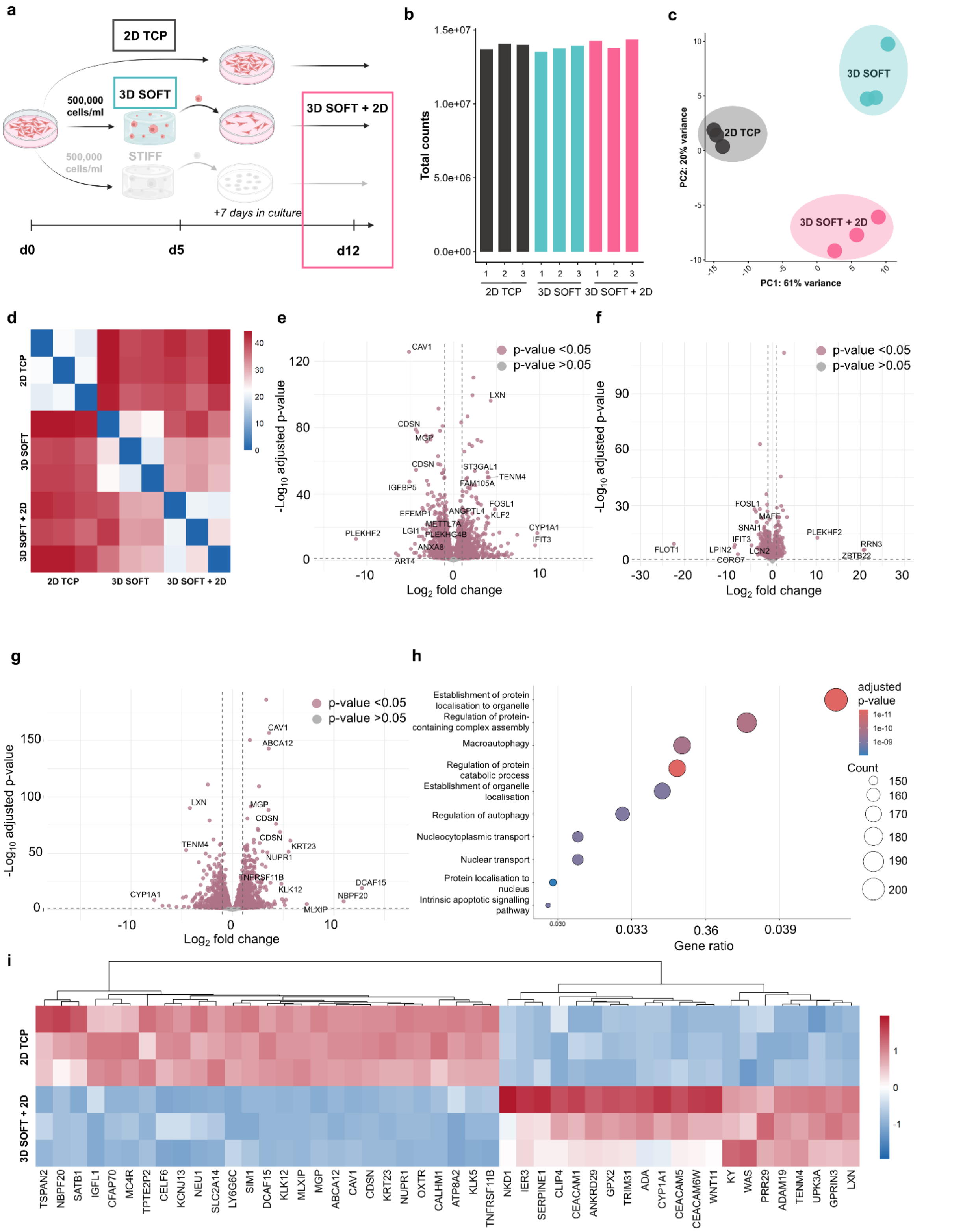


**Supplementary Fig. 7 | RNA-seq analyses in MCF-7 cells on 2D TCP, 3D soft alginate and retrieved and post-retrieval cells. a,** Graphical illustration of the experimental workflow. Schematics created with BioRender. **b,** RNA-seq library size per samples showing total read counts in cells on 2D TCP, 3D soft alginate and retrieved cells. **c,** Principal-component analysis (PCA) of variance-stabilised transformed (VST) gene expression profiles between groups. **d,** Sample-to-sample distance matrix based VST normalised counts, showing global transcriptional similarity and clustering structure. **e-f,** Volcano plots of DEGs (e) between 2D TCP and 3D soft alginate (n = 14821 genes), (f) between 3D soft alginate and retrieved cells (n =14000 genes) and **g,** between 2D TCP and retrieved cells (grey = not significant, salmon = adjusted p-value < 0.05, n = 14546 genes). **h,** GO biological process enrichment of DEGs (adjusted p-value < 0.1) identified by over-representation analysis. **i,** Heatmap of the top 50 DEGs ranked by absolute log2 fold change (VST) between 2D TCP and retrieved cells.

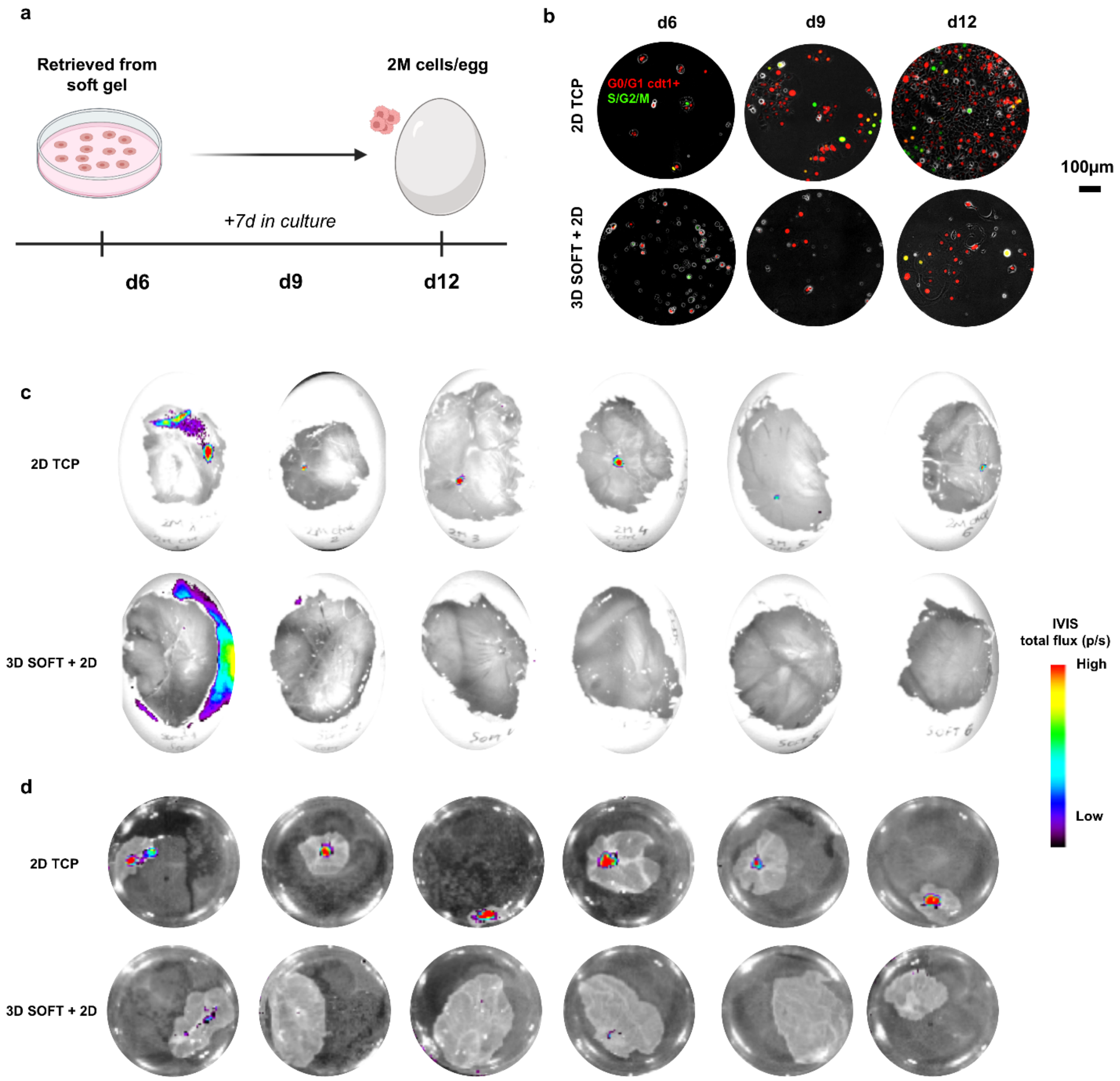


**Supplementary Fig. 8 | Tumour nodule formation in CAM models. a,** Graphical illustration of experimental workflow. Schematics created with BioRender. **b,** Representative fluorescence images of MCF-7 FUCCI2 cells ($G_0/G_1$ cdt1+ = mCherry-hCdt1 and $S/G_2/M$ = mVenus-hGeminin) during 2D monitoring at days 6, 9 and 12. **c-d,** IVIS images of MCF-7 FUCCI2 cells in CAM assay at day 19, 7 days after cell injection (n = 6 eggs per group), showing (c) in ovo and (d) ex ovo IVIS.

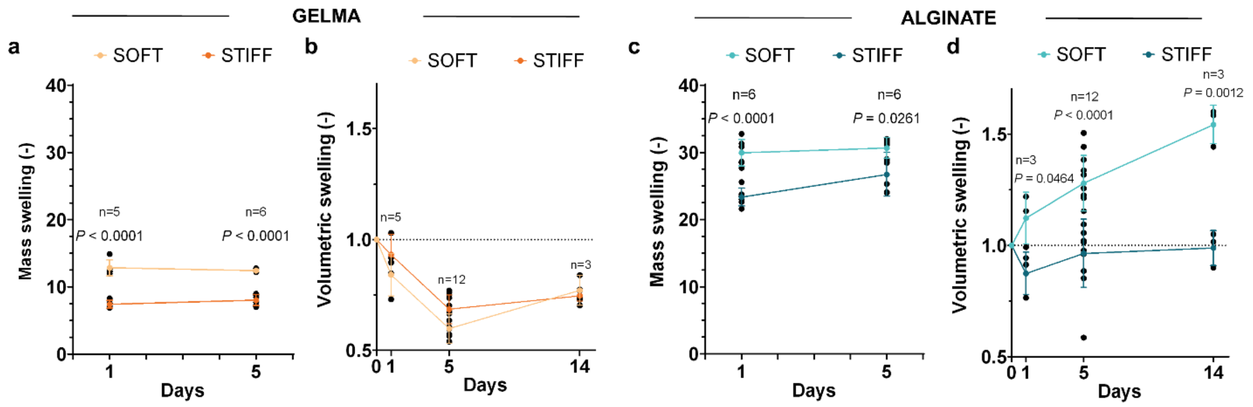


**Supplementary Fig. 9 | Mass and volumetric swelling characterisation of GelMA and alginate hydrogels over time a,** Mass swelling of soft and stiff GelMA hydrogels at days 1 and 5 with MCF-7 cells (n = 6). **b,** Volumetric swelling of soft and stiff GelMA hydrogels at days 1, 5 and 14 with MCF-7 cells (n = 3 – 12) (Volumetric swelling factors ($S_f$) of GelMA hydrogels at day 5 = 0.607 (soft) and 0.651 (stiff)). **c,** Mass swelling of soft and stiff alginate hydrogels at days 1 and 5 with MCF-7 cells (n = 6). **d,** Volumetric swelling of soft and stiff alginate hydrogels at days 1, 5 and 14 with MCF-7 cells (n = 3 – 12) (Volumetric swelling factors ($S_f$) of alginate hydrogels at day 5 = 1.258 (soft) and 0.929 (stiff)). Bars and error bars show mean ± s.d., respectively. Statistical significance (two-tailed Student's t-test (parametric)). a.u. stands for arbitrary units.

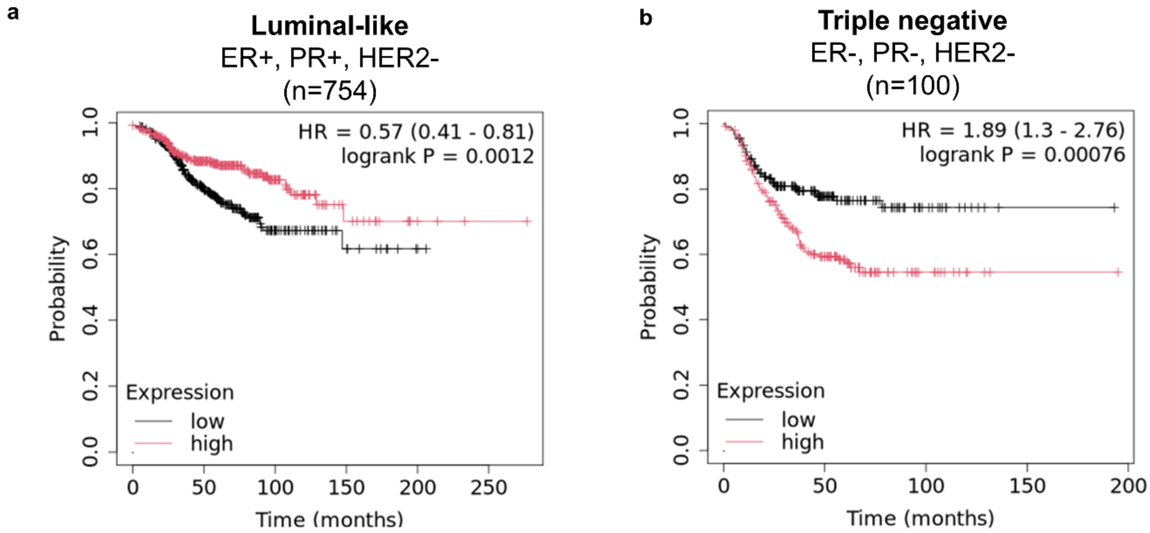


**Supplementary Fig. 10 | Distinct relapse-free survival of patients for different breast cancer subtypes.** Kaplan-Meier relapse-free survival plots according to FHL2 protein and mRNA expression levels for **a,** luminal and **b,** TNBC patients (n = number of patients). Data obtained from KMplot.com [1].

| CLINICAL DATA | BAG-DO-15 | HUB2322-020 | HD2322-085 | HUB2322-021 |
|---|---|---|---|---|
| **Hospital** | HU Donostia | HU Basurto | HU Donostia | HU Basurto |
| **Sex** | Female | Female | Female | Female |
| **Age at diagnosis** | 44 | 82 | 54 | 28 |
| **Menopausal status** | Premenopausal | Postmenopausal | Premenopausal | Premenopausal |
| **Family history** | Not specified | No | Yes | Yes |
| **Clinical stage at diagnosis** | III (localised) | - | II (localised) | |
| **ER (%)** | 100 | 95 | 99 | 95 |
| **PR (%)** | 85 | 5 (equivocal) | 99 | 90 |
| **HER2 score** | 1+ | 1+ | 2+ | 3+ |
| **HER2 ISH amplification** | Not applicable | Not applicable | Not amplified | Not applicable |
| **HER2 status** | Negative | Negative | Negative | Positive |
| **Ki67 (%)** | 40 | 40 | 25 | 40 |
| **Histological grade** | 3 | 2 | 2 | 2 |
| **Surgery date** | 2026/03/27 | - | 2026/03/18 | - |
| **Breast surgery** | Mastectomy | - | Lumpectomy | - |
| **Sample type** | Biopsy | Biopsy | Biopsy | Biopsy |
| **Anatomical location** | Breast | Right breast | Right breast | Left breast |
| **Breast biopsy classification** | B5a (invasive carcinoma) | - | - | - |
| **CK19** | Positive | Positive | Positive | Positive |

**Supplementary Table. 1 |** Pathological data from a luminal BC patient group (n=4).

| Reagent | Acro | Supplier, #Code | Stock information |
|---|---|---|---|
| High guluronic acid sodium alginate | MVG | Sigma-Aldrich, #42000101 | - |
| 2-(N-morpholino)ethanesulfonic acid | MES | Sigma-Aldrich, #M5287 | - |
| Sodium Chloride | NaCl | PanReac, #131659.1210 | - |
| N-hydroxysulfosuccinimide | NHS | Sigma-Aldrich, #130672 | - |
| N-(3-Dimethylaminopropyl)-N'-ethylcarbodiimide hydrochloride | EDC | Sigma-Aldrich, #E7750 | - |
| 5-norbornene-2-methylamine | N | TCI Chemicals, #N0907 | - |
| Hydroxylamine hydrochloride | HA · HCl | Sigma-Aldrich, #26103 | - |
| Spectra/Por 6 3.5 kDa MWCO prewetted kit | - | Spectrum, #132594 | - |
| 10X phosphate-buffered saline | 10X PBS | Gibco, #14200059 | - |
| Activated charcoal | - | Sigma-Aldrich, #C9157 | - |
| 0.22 µm Steritop-GP | - | Sigma-Aldrich, #S2GPT05RE | - |
| 0.22 µm Steriflip-GP | - | Sigma-Aldrich, #SCGP00525 | - |
| Deuterium oxide | D2O | Sigma-Aldrich, #1034280009 | - |
| Gelatine | - | Sigma-Aldrich, #G2500 | Porcine Type A, 300 bloom |
| DL-dithiothreitol | DTT | Sigma-Aldrich, #D0632 | Powdered |
| 1X phosphate-buffered saline | PBS | Gibco, #14190-094 | - |
| Lithium phenyl-2,4,6-trimethylbenzoylphosphinate | LAP | Sigma-Aldrich, #900889 | Powdered |
| Gelatin methacrylamide stock solution | GelMA | Gelomics, #SKU0010 | Porcine Type A, 300 bloom |
| Positive-displacement pipettes | - | Mettler-Toledo, #17008579 | - |
| Casting glass plate | - | 159 Filafarm, #1000185 | Borosilicate glass |
| Glass coating | Sigmacote | Sigma-Aldrich, #SL2 | - |
| 24-well plates | - | Thermo Fisher, #130186 | - |
| Low glucose Dulbecco's Modified Eagle's Medium | DMEM | Sigma-Aldrich, #D6046) | - |
| Fetal bovine serum | FBS | Gibco, #A5256701 | - |
| Penicillin/streptomycin | P/S | Gibco, #15140-122 | - |
| L-glutamine | - | Gibco, #25030-024 | - |

| | | | |
|---|---|---|---|
| PCR-based mycoplasma detection assay | - | Minerva Biolabs, #11-8100 | - |
| 0.05 % trypsin-EDTA | - | Gibco, #25300-054 | - |
| 0.5 % trypsin-EDTA | - | Gibco, #15400054 | - |
| 40 µm cell strainer | - | Thermo Fisher, #22-363-547 | - |
| 100 µm cell strainer | - | Corning, #CLS431752 | - |
| Calcein AM | - | ATT Bioquest #22002 | 4 mM stock solution in dimethyl sulfoxide (DMSO) |
| Ethidium homodimer-1 | EthD-1 | Sigma-Aldrich, #E1903 | 2 mM stock solution |
| Formaldehyde | - | Sigma-Aldrich, #HT501320 | - |
| Bovine serum albumin | BSA | Sigma-Aldrich, #A7906 | - |
| Triton X 100 | - | Sigma-Aldrich, #T8787 | - |
| Phalloidin Atto-633 | - | Sigma-Aldrich, #68825 | 20 µM stock solution in methanol |
| Phalloidin Atto-550 | - | Sigma-Aldrich, #19083 | 20 µM stock solution in methanol |
| Phalloidin Atto-488 | - | Sigma-Aldrich, #49409 | 20 µM stock solution in Milli-Q water |
| 4′,6-diamidino-2-phenylindole | DAPI | Sigma-Aldrich, #MBD0015 | 1 mg/mL stock solution |
| 3D CellTiter-Glo | - | Promega, #G9683 | - |
| White 96-well plates | - | Corning, #3903 | - |
| Alginate lyase | - | Sigma-Aldrich, #A1603 | Powdered |
| 10X Hank's Balanced Salt Solution | HBSS | Gibco, #11570476 | - |
| Maxwell RSC simplyRNA Cells Kit | - | Promega, #AS1390 | - |
| Maxima First-Strand cDNA synthesis kit | - | Thermo Fisher, #15273796 | - |
| TB Green Premix Ex Taq | - | Takara Bio, #RR820W | - |
| Lipofectamine 2000 | - | Invitrogen, #11668030 | - |
| 6-well plate | - | Corning, #3506 | - |
| Electronic diamond scriber | - | Medite Medical, #01-2000-00 | EDS 20 |
| Matrigel | - | Corning, #356237 | - |
| MACS Tissue Storage Solution | - | MACS Milteny, #130-100-008 | - |
| Sterile disposable blades | - | Swann_morton, #ABH5011 | - |

| | | | |
|---|---|---|---|
| 10x collagenase/hyaluronidase | - | STEMCELL, #07912 | - |
| Collagenase Type II | - | Gibco, #17101015 | - |
| Advanced DMEM-F12 | - | Gibco, #12634010 | - |
| 4-(2-hydroxyethyl)-1-piperazineethanesulfonic acid | HEPE S | Gibco, #15630080 | - |
| Glutamax | - | Gibco, #35050061 | - |
| ACK Lysing Buffer | - | Gibco, #A1049201 | - |
| Cryostor CS-10 freezing media | - | STEMCELL, #07930 | - |
| β-estradiol | - | Sigma-Aldrich, #E2257 | 100 µM stock solution in 100 % ethanol |
| Tamoxifen | - | Selleck Chem., #S1238 | 10mM stock solution in DMSO |
| Dimethyl sulfoxide | DMS O | Sigma-Aldrich, #D5879 | - |
| Glass coverslips | - | Epredia #24X50-1-002G | - |
| Qubit RNA HS Assay Kit | - | Thermo Fisher, #Q32855 | - |
| Agilent RNA 6000 Nano Chips | - | Agilent Tech., #5067-1511 | - |
| Illumina Stranded mRNA Prep kit | - | Illumina, #20040532 | - |
| Qubit dsDNA HS Assay Kit | - | Thermo Fisher, #Q32854 | - |
| Agilent High Sensitivity DNA Kit | - | Agilent Tech., #5067-4626 | - |
| Nunc™ Non-Treated 24-well plate | - | Thermo Fisher, #144530 | - |
| Growth Factor Reduced (GFR) Basement Membrane Matrix, Phenol Red-free, LDEV-free | - | Corning, 356231 | - |

**Supplementary Table. 2 |** Materials and reagents.

| Output | Name | Formula | Parameters |
|---|---|---|---|
| $DS_{real}$ (%) | Degree of substitution for GelMA | $1 - [\text{Area of lysine methylene (GelMA)} / \text{Area of lysin methylene (Gelatine)}] \cdot 100$ | |
| E | Elastic Modulus | $2 \cdot G^{*} \cdot (1 + \vartheta)$ | $\vartheta$ the Poisson's ratio, a value of 0.5 for hydrogels [2] |
| G* | Complex Modulus | $\sqrt{(G')^2 + (G")^2}$ | G' and G'' are storage and loss moduli, respectively |
| $\xi$ (ref. [3]) | Mesh size | $\sqrt[3]{6RT/G'\pi N_{av}}$ | $N_{av}$ Avogadro's number ($6.022 \cdot 10^{23}$ 1/mol), R the ideal gas constant (8.314 $m^3$Pa/molK) and T the RT (293 K) |
| $S_t$ | Volumetric swelling | $\frac{V_t}{V_0}$ | $V_t$ refers to the hydrogel volume at a given time point t, and $V_0$ at the initial time point |
| $Q_m$ | Mass swelling | $\frac{(W_w - W_d)}{W_d}$ | $W_w$ refers to the fully hydrated weight and $W_d$ is the dry weight after lyophilisation |

**Supplementary Table. 3 |** Theoretical formulas.

| **Antibody** | **Supplier and product code** | **Stock concentration** | **Dilution** |
|---|---|---|---|
| anti-E-cadherin | Cell Signaling Technology, #3195S | 54 µg/mL | 1:100 |
| anti-Vimentin | abcam, #ab8069 | 1000 µg/mL | 1:500 |
| anti-Ki67 | Invitrogen, #14-5698-82 | 500 µg/mL | 1:250 |
| anti-p27$^{kip1}$ | abcam, #ab32034 | 566 µg/mL | 1:500 |
| anti-phospho-p38 | Cell Signaling Technology, #9216S | 479 µg/mL | 1:400 |
| anti-phospho-p44/42 MAPK (p-ERK1/2) | Cell Signaling Technology, #9101S | 191 µg/mL | 1:400 |
| anti-SOX9 | abcam, #ab185230 | 1026 µg/mL | 1:200 |
| anti-COUP TF1 | abcam, #ab181137 | 566 µg/mL | 1:250 |
| anti-YAP | Santa Cruz Biotechnology, #sc-101199 | 100 µg/mL | 1:100 |
| anti-FHL2 | Sigma-Aldrich, #HPA006028 | 100 µg/mL | 1:100 |
| anti-PanCK | Invitrogen, # MA5-13203 | 200 µg/mL | 1:100 |
| Alexa Fluor 488 | Cell Signaling Technology, #4412S | 2000 µg/mL | 1:500 |
| Alexa Fluor 555 | Cell Signaling Technology, #4417 | 2000 µg/mL | 1:500 |
| Alexa Fluor 647 | Cell Signaling Technology, #4410S | 2000 µg/mL | 1:500 |

**Supplementary Table. 4 |** Primary and secondary antibodies.

| Gene | Forward | Reverse |
|---|---|---|
| *Cdkn1a* | TGTCTTGTACCCTTGTGCCT | AATCTGTCATGCTGGTCTGC |
| *Tp53* | CCTCCTCAGCATCTTATCCGA | GGTACAGTCAGAGCCAACCT |
| *Bhlhe41* | AAGGAGCATGAAACGAGACGA | CTCGGTTAAGGCGGTTAAAGC |
| *Sox9* | CTCGCCACACTCCTCCTC | CTGCACGTCGGTTTTGGG |
| *Nr2f1* | ATCGTGCTGTTCACGTCAGAC | TGGCTCCTCACGTACTCCTC |
| *Fhl2* | GGAGTTGGGGAGACTGGTTG | TTGGCGAACAGGGTCTCAAA |
| *Yap1* | AGGTTGGGAGATGGCAAAGA | CATCATATTCTGCTGCACTGGT |
| *Gapdh* | CACATCGCTCAGACACCATG | TTGAGGTCAATGAAGGGGTC |
| *Pcbp1* (ref. [4]) | TGATCATCGACAAGCTGGAG | TCTTTGATCTTACACCCGCC |
| *Ccser2* (ref. [4]) | GACAGGAGCATTACCACCTCAG | CTTCTGAGCCTGGAAAAAGGGC |

**Supplementary Table. 5 |** Forward and reverse primer sequences of dormancy genes analysed by RT-qPCR.

| 3D organoids<br>HD085TR1 | | | |
|---|---|---|---|
| **Complement** | **Concentration** | **Supplier** | **Reference** |
| Oncopro Medium | 1x | Gibco | #A5701201 |
| Oncopro Supplement | 1x | Gibco | #A5701201 |
| Oncopro BSA | 1x | Gibco | #A5701201 |
| P/S | 2% | Gibco | #15140-122 |
| B27 | 1x | Gibco | #A5701201 |
| FGF-10 | 10ng/mL | PeproTech | #AF-100-26-25UG |
| Primocin | 100µg/mL | InvivoGen | #ant-pm-1 |
| β-estradiol | 10nM | Sigma-Aldrich | #E2257-1MG |
| Y-27632 | 10uM | AbMole | #M1817 |
| **3D organoids**<br>HUB020, HUB021 | | | |
| **Complement** | **Concentration** | **Supplier** | **Reference** |
| Mammocult media | 1x | STEMCELL | #05620 |
| MammoCult Supplementary | 1x | STEMCELL | #05620 |
| Hydrocortisone | 0.48 µg/mL | STEMCELL | #07925 |
| Heparin | 4 µg/mL | STEMCELL | #07980 |
| P/S | 1% (v/v) | Gibco | #15140-122 |
| Primocin | 100 µg/mL | InvivoGen | ant-pm-1 |
| **2D culture** (adapted from Bock et al 2023 [5])<br>BAG-DO-15 | | | |
| **Complement** | **Concentration** | **Supplier** | **Reference** |
| Advanced DMEM-F12 | 1x | Gibco | #12634010 |
| HEPES | 1% (v/v) | Gibco | #15630080 |
| P/S | 1% (v/v) | Gibco | #15140-122 |
| Glutamax | 1% (v/v) | Gibco | #35050061 |
| EGF | 10 ng/mL | PeproTech | #AF-100-15-100UG |
| Y-27632 | 5 µM | MedChemExpress | HY-10583 |
| Hydrocortisone | 1 µg/mL | Sigma-Aldrich | #H088-1G |
| Primocin | 100 µg/mL | InvivoGen | #ant-pm-05 |
| FBS | 10% | Gibco | #A5256701 |

**Supplementary Table. 6 |** Culture conditions and media compositions of different group of patients.

| Marker | Channel | Illumination wavelength | LED intensity | Exposure time | Filter |
|---|---|---|---|---|---|
| S/G2/M - mVenus-hGeminin | phYFP | 511 nm | 100 % | 200 ms | 46 HE |
| G0/G1 cdt1+ - mCherry-hCdt1 | mCherry | 555 nm | 100 % | 200 ms | 45 |
| Live cells | Calcein AM | 469 nm | 4 % | 10 ms | 38 HE |
| Dead cells | EthD-1 | 555 nm | 50 % | 50 ms | 45 |
| Ki67 | Alexa Fluor 555 | 555 nm | 80 % | 100 ms | 45 |
| p27 | Alexa Fluor 488 | 469 nm | 20 % | 30 ms | 38 HE |
| Phalloidin green | Atto-488 | 469 nm | 20 % | 30 ms | 38 HE |
| Phalloidin red | Atto-633 | 632 nm | 20 % | 20 ms | 50 |
| DAPI | DAPI | 385 nm | 4 % | 10 ms | 96 HE |

**Supplementary Table. 7 |** Zeiss AxioObserver 7 epifluorescence image acquisition data for 3x3 binning.

| **Marker** | **Channel** | **Excitation wavelength** | **Detection wavelength** | **Laser intensity** | **Detector gain** |
|---|---|---|---|---|---|
| S/G2/M - mVenus-hGeminin | phYFP | 524 nm | 400 / 580 nm | 4.50 % | 800 V |
| G0/G1 cdt1+ - mCherry-hCdt1 | mCherry | 587 nm | 400 / 635 nm | 4.50 % | 750 V |
| SOX9 | Alexa Fluor 488 | 493 nm | 490 / 700 nm | 1 % | 500 V |
| NR2F1 | Alexa Fluor 488 | 493 nm | 490 / 700 nm | 1 % | 500 V |
| p27 | Alexa Fluor 488 | 493 nm | 490 / 700 nm | 1 % | 500 V |
| FHL2 | Alexa Fluor 488 | 493 nm | 490 / 700 nm | 1 % | 500 V |
| P-ERK | Alexa Fluor 488 | 493 nm | 400 / 555 nm | 1 % | 700 V |
| Ki67 | Alexa Fluor 555 | 553 nm | 540 / 700 nm | 2 % | 700 V |
| YAP | Alexa Fluor 647 | 653 nm | 645 / 700 nm | 1 % | 700 V |
| P-p38 | Alexa Fluor 647 | 653 nm | 645 / 700 nm | 1 % | 700 V |
| PanCK | Alexa Fluor 647 | 653 nm | 645 / 700 nm | 0.5 % | 700 V |
| Phalloidin green | Atto-488 | 653 nm | 645 / 700 nm | 1 % | 700 V |
| Phalloidin orange | Atto-550 | 553 nm | 545 / 617 nm | 1 % | 600 V |
| Phalloidin red | Atto-633 | 631 nm | 645 / 700 nm | 0.5 % | 700 V |
| DAPI | DAPI | 353 nm | 400 / 560 nm | 1 % | 700 V |

**Supplementary Table. 8 |** Zeiss LSM 900 confocal image acquisition data with 1 airy unit pinhole aperture.
